# Convergent evolution of linked mating-type loci in basidiomycete fungi

**DOI:** 10.1101/626911

**Authors:** Sheng Sun, Marco A. Coelho, Joseph Heitman, Minou Nowrousian

## Abstract

Sexual development is a key evolutionary innovation of eukaryotes. In many species, mating involves interaction between compatible mating partners that can undergo cell and nuclear fusion and subsequent steps of development including meiosis. Mating compatibility in fungi is governed by mating type determinants, which are localized at mating type (*MAT*) loci. In basidiomycetes, the ancestral state is hypothesized to be tetrapolar (bifactorial), with two genetically unlinked *MAT* loci containing homeodomain transcription factor genes (*HD* locus) and pheromone and pheromone receptor genes (*P/R* locus), respectively. Alleles at both loci must differ between mating partners for completion of sexual development. However, there are also basidiomycete species with bipolar (unifactorial) mating systems, which can arise through genomic linkage of the *HD* and *P/R* loci. In the order *Tremellales*, which is comprised of mostly yeast-like species, bipolarity is found only in the human pathogenic *Cryptococcus* species. Here, we describe the analysis of *MAT* loci from the *Trichosporonales*, a sister order to the *Tremellales*. We analyzed genome sequences from 29 strains that belong to 24 species, including two new genome sequences generated in this study. Interestingly, in all of the species analyzed, the *MAT* loci are fused and a single *HD* gene is present in each mating type. This is similar to the organization in the pathogenic Cryptococci, which also have linked *MAT* loci and carry only one *HD* gene per *MAT* locus instead of the usual two *HD* genes found in the vast majority of basidiomycetes. However, the *HD* and *P/R* allele combinations in the *Trichosporonales* are different from those in the pathogenic *Cryptococcus* species. The differences in allele combinations compared to the bipolar Cryptococci as well as the existence of tetrapolar *Tremellales* sister species suggest that fusion of the *HD* and *P/R* loci and differential loss of one of the two *HD* genes per *MAT* allele occurred independently in the *Trichosporonales* and pathogenic Cryptococci. This finding supports the hypothesis of convergent evolution at the molecular level towards fused mating-type regions in fungi, similar to previous findings in other fungal groups. Unlike the fused *MAT* loci in several other basidiomycete lineages though, the gene content and gene order within the fused *MAT* loci are highly conserved in the *Trichosporonales*, and there is no apparent suppression of recombination extending from the *MAT* loci to adjacent chromosomal regions, suggesting different mechanisms for the evolution of physically linked *MAT* loci in these groups.

**Author summary:** Sexual development in fungi is governed by genes located within a single mating type (*MAT*) locus or at two unlinked *MAT* loci. While the latter is thought to be the ancestral state in basidiomycetes, physical linkage of the two *MAT* loci has occurred multiple times during basidiomycete evolution. Here, we show that physically linked *MAT* loci are present in all analyzed species of the basidiomycete order *Trichosporonales*. In contrast to previously studied basidiomycetes, the fused *MAT* loci in the *Trichosporonales* have highly conserved gene order, suggesting that this fusion might date back to the common ancestor of this lineage.

## Introduction

Sexual reproduction is prevalent among eukaryotic organisms, but despite its rather conserved core features (syngamy/karyogamy and meiosis), many aspects of sexual development show high evolutionary flexibility [1-3]. This includes the determination of compatible mating partners that can successfully undergo mating and complete the sexual cycle. In many species, compatibility is determined by one or more genetic loci that differ between compatible mating partners. The evolution of such genes has been studied in many systems including plants, animals, algae, and fungi [1, 4-6]. In animals and plants, genes that determine sexual compatibility are often found on sex chromosomes, which have distinct patterns of inheritance and diversification compared to autosomes [7]. In fungi, genetic systems for determining mating compatibility vary widely, ranging from single genetic loci on autosomes to chromosomes that show suppressed recombination over most of the chromosome, similar to sex chromosomes. Studies in different fungal groups have revealed that transitions towards larger, sex chromosome-like regions have occurred multiple times in fungal evolution, with some systems having evolved only recently [8-10]. Thus, fungi are excellent model systems to study the evolution of genomic regions involved in mating and mating type determination.

Mating compatibility in fungi is governed by mating-type genes, which are located in mating-type (*MAT*) loci [11, 12]. While ascomycetes and the Mucoromycotina are bipolar (unifactorial), i.e. harboring only one *MAT* locus [13-15], in basidiomycetes, the ancestral state is considered to be tetrapolar (bifactorial), with two genetically unlinked *MAT* loci controlling mating-type determination at the haploid stage [16, 17]. One *MAT* locus usually contains tightly linked pheromone and pheromone receptor genes (*P/R* locus) that are involved in pre-mating recognition, and the other (*HD* locus) harbors homeodomain transcription factor genes encoding homeodomain (HD) proteins of class 1 and class 2, which determine viability after syngamy. Importantly, alleles at both loci must differ between mating partners for completion of the sexual cycle [18].

However, there are also many basidiomycete species with bipolar mating systems, which can arise through genomic linkage of the *HD* and *P/R* loci, or when the *P/R* locus loses its function in determining mating specificity [16, 18, 19]. Bipolarity through *MAT* loci fusion is found in the subphylum *Ustilaginomycotina* in *Malassezia* species and *Ustilago hordei*, several *Microbotryum* species (subphylum *Pucciniomycotina*), as well as in the pathogenic Cryptococci from the class *Tremellomycetes* (subphylum *Agaricomycotina*) [1, 8, 20-24] (Fig. 1). In *Microbotryum*, several convergent transitions to linked *MAT* loci have occurred within the genus [8]. One shared feature of known species with fused (physically linked) *MAT* loci is that they are associated with plant or animal hosts as pathogens or commensals. It has been hypothesized that the necessity to find a mating partner while associated with a host might have favored linkage of *MAT* loci, because having one linked instead of two unlinked *MAT* loci increases the compatibility among siblings that are descendants from a single pair of parents. This would improve mating compatibility rates on a host where other mating partners might be difficult to find, for instance when a host is initially colonized by a single diploid genotype or meiotic progeny from a single tetrad [4, 9, 16, 18, 25].

**Fig 1.**
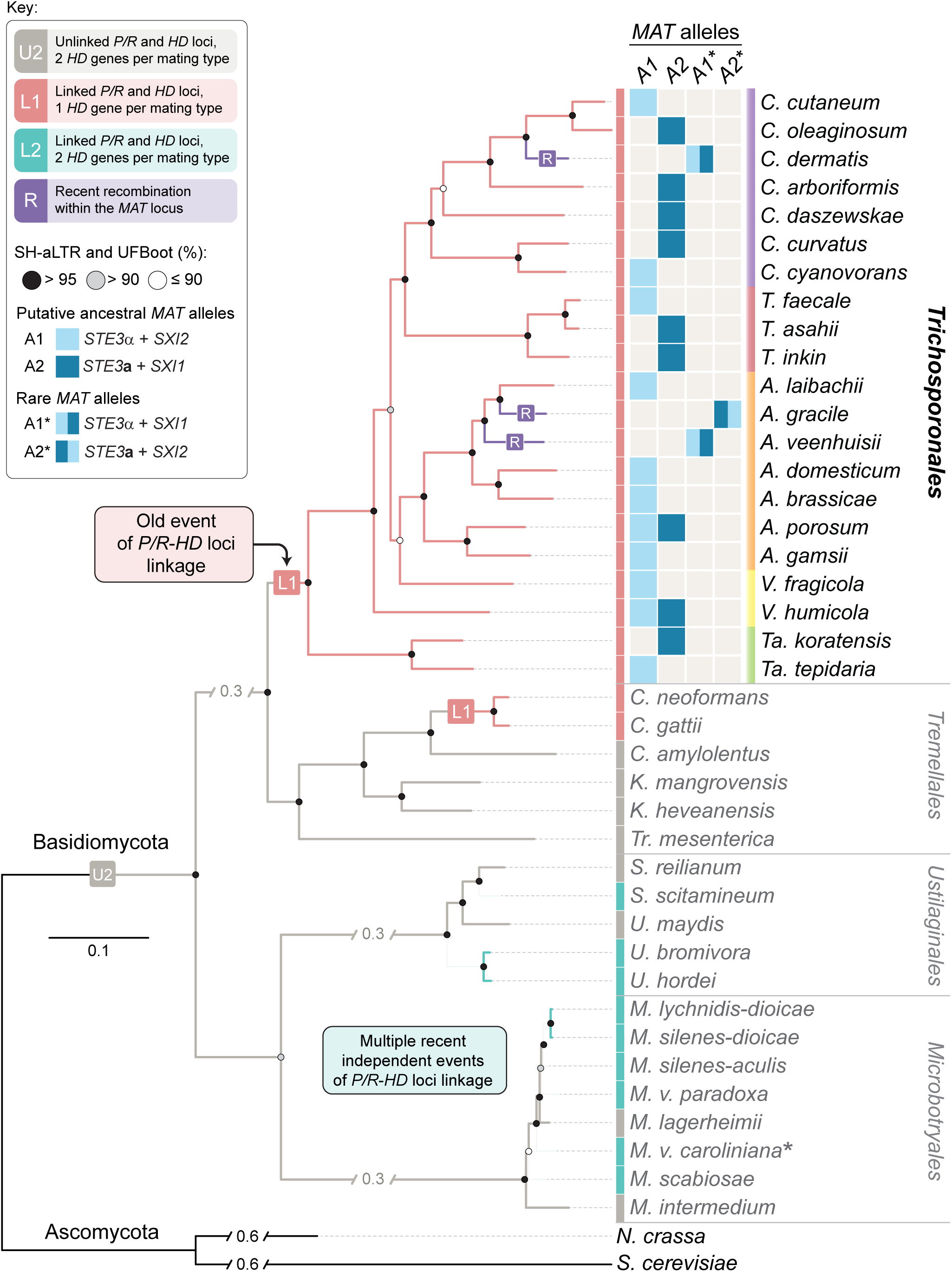
Overview of the haploid *Trichosporonales* species for which *MAT* regions were analyzed. Shown here is a phylogenetic tree of basidiomycetes with a focus on groups where fusion of *MAT* loci occurred as was shown previously (*Cryptococcus, Ustilago, Malassezia, Microbotryum, Sporisorium*) [4, 8, 18]. The phylogenetic branching within the tree agrees with previous studies [4, 8, 40, 51, 141]. Next to each *Trichosporonales* species name (color coded according to their genus), the *MAT* alleles that are represented in the sequenced genomes are shown. Within one species, different strains carrying different alleles are currently known only for *V. humicola* and *A. porosum*. Abbreviations for *Trichosporonales* genus names are: *A – Apiotrichum, C – Cutaneotrichosporon, T – Trichosporon, Ta – Takashimella, V – Vanrija*. Abbreviations for other basidiomycete genus names are: *C – Cryptococcus, K – Kwoniella, M – Microbotryum, S – Sporisorium, Tr – Tremella, U – Ustilago*. Ascomycetes *Neurospora crassa* and *Saccharomyces cerevisiae* were used as outgroups. Branch lengths are given in number of substitutions per site (scale bar). Branch support was assessed by 10,000 replicates of both ultrafast bootstrap approximation (UFBoot) and the Shimodaira-Hasegawa approximate likelihood ratio test (SH-aLRT) and represented as circles colored as given in the key. The asterisk next to the species name for *Microbotryum violaceum caroliniana* indicates that the *HD1* gene was specifically lost in the A2 mating type of this species [8].

Two additional evolutionary features can be associated with the linkage and expansion of the *MAT* loci. One is the presence of other development-associated genes in the fused *MAT* loci, and the second is suppression of recombination at the fused *MAT* loci that can extend along the *MAT*-containing chromosome [12, 24, 26]. Suppression of recombination is a hallmark of sex chromosomes in other eukaryotes, and thus might point towards convergent evolutionary transitions for the regulation of sexual development in eukaryotes [27, 28]. The evolution of recombination suppression after the physical linkage of *MAT* loci would further increase compatibility under inbreeding conditions, and recruitment of sex-associated genes into the *MAT* locus might facilitate the inheritance of favorable allele combinations through genetic linkage [29].

An instance of physically linked *MAT* loci has been well-studied in *Cryptococcus neoformans*, a member of a group of closely related, pathogenic *Cryptococcus* species [1]. Fusion of the *HD* and *P/R* loci most likely occurred in the ancestor of the pathogenic Cryptococci, because other analyzed *Tremellales* species including the closely related, but non-pathogenic *Cryptococcus amylolentus, Kwoniella heveanensis, Kwoniella mangrovensis, Cryptococcus wingfieldii*, and *Cryptococcus floricola* are all tetrapolar [30-34]. The *C. neoformans MAT* locus, which consists of genetically linked *HD* and *P/R* loci, encompasses more than 20 genes over a region spanning more than 100 kb and has two alleles designated **a** and α. In the majority of basidiomycetes, each *MAT* allele at the *HD* locus carries both the *HD1* and the *HD2* transcription factor genes, whereas in *C. neoformans*, the *MAT*α allele contains only the *HD1*-class gene *SXI1*α, and *MAT***a** contains only the *HD2*-class gene *SXI2***a**. Except for a gene conversion hotspot, the *C. neoformans MAT* locus displays suppressed meiotic recombination [24, 35, 36].

Among the *Tremellomycetes*, pathogenic Cryptococci are the only species for which fused *MAT* loci have been described so far [18, 24]. In a previous study, we analyzed the genome sequence of *Cutaneotrichosporon oleaginosum* strain IBC0246 (formerly *Trichosporon oleaginosus*), which belongs to the *Trichosporonales*, a sister order to the *Tremellales* within the *Tremellomycetes* class. *Trichosporonales* species are widely distributed in the environment and have been isolated from a variety of substrates including soil, decaying plant material, and water. Many species are saprobes, but some have also been found to be associated with animals including humans either as commensals or pathogens [37-39]. Despite their common occurrence in the environment, *Trichosporonales* are an understudied fungal group, and sexual reproduction has not yet been observed for any of the known species [40, 41]. Recently, several *Trichosporonales* were studied with respect to their biotechnological properties, including the oil-accumulating *C. oleaginosum*, which was first isolated from a dairy plant, and has the ability to metabolize chitin-rich and other non-conventional substrates [42-45]. The sequenced *C. oleaginosum* strain is haploid and similar in genome size and gene content to strains from the sister order *Tremellales*; this was also the case for several other *Trichosporonales* genomes that have since been sequenced [44, 46-51]. Interestingly, *C. oleaginosum* showed some similarities to *C. neoformans* in the organization of *MAT* loci. This included the presence of several genes with functions during mating into the *HD* and *P/R* loci, as well as the presence of only a *SXI1* homolog at the *HD* locus [44]. However, in the draft genome assembly, *HD* and *P/R* loci were situated on different scaffolds. Furthermore, a sexual cycle has not yet been described for *C. oleaginosum*. Thus, it was not possible to conclusively distinguish whether this species carries two unlinked or fused *MAT* loci, although it seemed more likely that it utilized a tetrapolar mating system as this is present in the majority of species from the sister group of *Tremellales*.

Since the analysis of the first *C. oleaginosum* genome, several more *Trichosporonales* genomes have been sequenced, although none were analyzed with respect to their *MAT* loci [46-52]. Here, we describe the sequencing of two additional *Trichosporonales* genomes for *C. oleaginosum* ATCC20508 and *Vanrija humicola* CBS4282, and the analysis of *MAT* loci organization in *Trichosporonales* genomes from 24 different species. Interestingly, we found that all of the species analyzed have physically linked *MAT* loci that contain both *P/R* and *HD* genes. Furthermore, all the analyzed strains of each species contain only one of the two ancestral *HD* genes (*SXI1* and *SXI2*), with almost all of the species carrying *HD* and *P/R* allele combinations that are different from those found at the *MAT* loci of pathogenic Cryptococci [35]. The differences in allele combinations as well as the existence of tetrapolar *Tremellales* sister species to the bipolar Cryptococci suggest that the fusion of the *HD* and *P/R* loci as well as the loss of one *HD* gene per allele occurred independently in the *Trichosporonales* and the pathogenic Cryptococci. This provides further evidence that convergent evolution leading to physically linked *MAT* loci is evolutionary beneficial under certain circumstances, and has been selected for multiple times in basidiomycetes. However, in contrast to other basidiomycetes, such as the pathogenic Cryptococci and several *Microbotryum* species [8, 35], the fused *MAT* loci of the *Trichosporonales* have a highly conserved gene content and gene order, suggesting unique mechanisms underlying the evolution of fused *MAT* loci in these groups.

## Results

### *Trichosporonales* species have fused *HD* and *P/R* mating type loci

Since 2015, genome sequences have been published for *Trichosporonales* species from the genera *Apiotrichum, Cutaneotrichosporon, Takashimella, Trichosporon*, and *Vanrija* [46-53] (Table 1). We analyzed the genomes of 29 isolates that belong to 24 *Trichosporonales* species for the organization of the *MAT* loci, and found that all of them contain fused *MAT* loci, i.e. the *MAT* loci are physically linked on the same contig, with the mating-type determining genes for the *P/R* locus (*STE3* and the pheromone precursor genes) and the *HD* locus (*SXI* homeodomain genes) located ∼55 kb apart from each other. The gene content between the *P/R* and *HD* loci is nearly identical in all species except for the early-branching genus *Takashimella* (Fig 2). Except for four strains described below, all analyzed *MAT* loci comprise either the combination of the *HD* gene *SXI1* with the pheromone allele *STE3***a**, or the combination of the *SXI2* gene and *STE3*α allele (Table 1, Figs 1 and 2). Each *MAT* allele carries a single pheromone precursor gene in the vicinity of the *STE3* gene (Fig. 2, S1 Text). The *SXI*/*STE3* combinations found in the majority of *Trichosporonales* are different from the *MAT* alleles in *C. neoformans*, where the *SXI1* gene is combined with *STE3*α in the *MAT*α allele, and *SXI2* is combined with *STE3***a** in the *MAT***a** allele [24, 35]. To avoid confusion with the *C. neoformans* nomenclature and to be compatible with allele designations in other basidiomycetes, we named the *STE3*α-containing *MAT* allele A1, and the *STE3***a**-containing *MAT* allele A2 (Figs 1 and 2), with each allele containing physically linked *HD* and *P/R* loci.

**Table 1.** Genome assemblies that were analyzed in this study.

| Genus | Species | Strain | MAT allele<sup>2</sup> | GenBank acc. number | Reference |
| --- | --- | --- | --- | --- | --- |
| *Apiotrichum* | *brassicae* | JCM1599 | A1 | BCJI00000000 | [51] |
| *Apiotrichum* | *domesticum* | JCM9580 | A1 | BCFW01000000 | [46] |
| *Apiotrichum* | *gamsii* | JCM9941 | A1 | BCJN00000000 | [51] |
| *Apiotrichum* | *gracile* | JCM10018 | A2* | BCJO00000000 | [51] |
| *Apiotrichum* | *laibachii* | JCM2947 | A1 | BCKV01000000 | [51] |
| *Apiotrichum* | *porosum* | DSM27194 | A1 | RSCE00000000 | [52] |
| *Apiotrichum* | *porosum* | JCM1458 | A2 | BCJG01000000 | [51] |
| *Apiotrichum* | *veenhuisii* | JCM10691 | A1* | BCKJ00000000 | [51] |
| *Cutaneotrichosporon* | *arboriformis* | JCM14201 | A2 | BEDW01000001 | [51] |
| *Cutaneotrichosporon* | *curvatus* | DSM101032 | A2 | LDEP01000000 | [48] |
| *Cutaneotrichosporon* | *cutaneum* | JCM1462 | A1 | BCKU01000000 | [51] |
| *Cutaneotrichosporon* | *cyanovorans* | JCM31833 | A1 | BEDZ01000001 | [51] |
| *Cutaneotrichosporon* | *daszewskae* | JCM11166 | A2 | BEDX01000001 | [51] |
| *Cutaneotrichosporon* | *dermatis* | JCM11170 | A1* | BCKR01000001 | [51] |
| *Cutaneotrichosporon* | *mucoides* | JCM9939<sup>1</sup> | A1*, A2* | BCJT01000001 | [51] |
| *Cutaneotrichosporon* | *oleaginosum* | ATCC20508 | A2 | QKWL01000000 | this study |
| *Cutaneotrichosporon* | *oleaginosum* | ATCC20509 | A2 | MATS01000000 | [47] |
| *Cutaneotrichosporon* | *oleaginosum* | IBC0246 | A2 | JZUH00000000 | [44] |
| *Takashimella* | *koratensis* | JCM12878 | A2 | BCKT01000001 | [51] |
| *Takashimella* | *tepidaria* | JCM11965 | A1 | BCKS01000001 | [51] |
| *Trichosporon* | *asahii* | CBS8904 | A2 | JH925096 | [140] |
| *Trichosporon* | *coremiiforme* | JCM2938<sup>1</sup> | A1, A2 | JXYL01000000 | [50] |
| *Trichosporon* | *faecale* | JCM2941 | A1 | JXYK01000000 | [50] |
| *Trichosporon* | *inkin* | JCM9195 | A2 | JXYM01000000 | [50] |
| *Trichosporon* | *ovoides* | JCM9940<sup>1</sup> | A1, A1 | JXYN01000000 | [50] |
| *Vanrija* | *fragicola* | JCM1530 | A1 | BEDY01000001 | [51] |
| *Vanrija* | *humicola* | CBS4282 | A1 | QKWK01000000 | this study |
| *Vanrija* | *humicola* | JCM1457 (CBS571) | A2 | BCJF01000000 | [51, 53] |
| *Vanrija* | *humicola* | JCM9575 (UJ1) | A2 | BFAH01000000 | [49] |

<sup>1</sup>Hybrid strain carrying two MAT alleles derived from the two different parental species that underwent hybridization (S2 Text).
<sup>2</sup>MAT alleles contain the following HD and P/R allele combinations: A1, *STE3α*+*SXI2*; A2, *STE3a*+*SXI1*; A1\*, *STE3α*+*SXI1*; A2\*, *STE3a*+*SXI2*.

**Fig 2.**
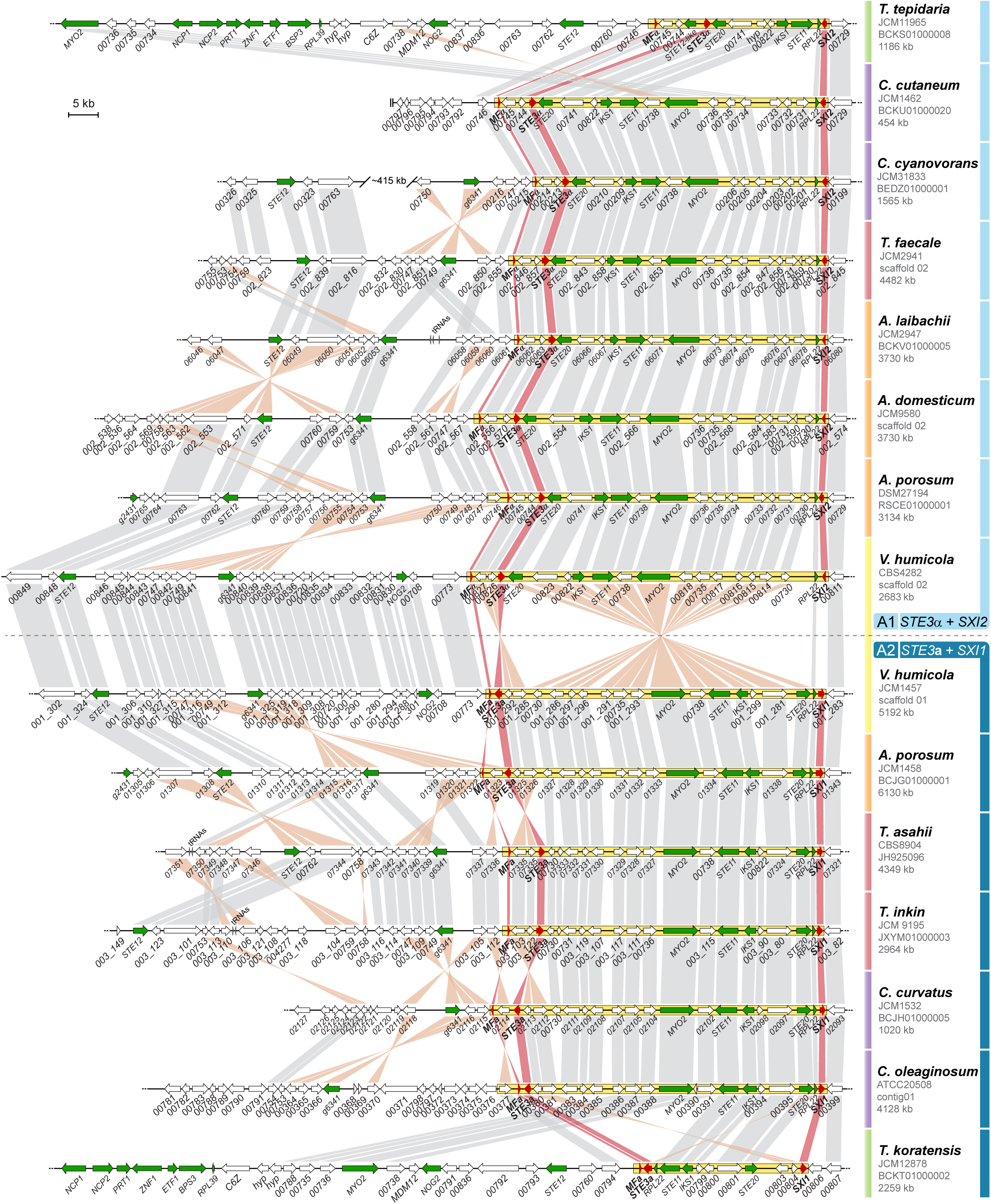
*MAT* loci of *Trichosporonales*. Classic mating type-defining genes (*SXI1* or *SXI2, STE3*, and the pheromone precursor genes) are shown in red, other genes that are part of the mating-type locus or flanking the mating-type locus (gene *g6341*) of *C. neoformans* are shown in green, and genes not present in the *C. neoformans* mating type locus are shown in white. The *MAT* allele is indicated on the right, with A1 and A2 *MAT* alleles shown above and below the dashed line, respectively. For simplicity, the A1* and A2* alleles are not included in this Figure, and are instead depicted in S5 and S6 Figs. The proposed *MAT* locus region is enclosed in a yellow box in each strain. Orthologs are connected by grey or orange bars when in the same or opposite orientations, respectively.

The fact that the genomes of 24 *Trichosporonales* species carry fused *MAT* loci with mostly identical gene content in two allelic combinations distributed throughout the phylogenetic tree (Fig 1, see also Fig 2 and the next results section) suggests that the event physically linking the two *MAT* loci predates the diversification of the *Trichosporonales* clade. However, at the beginning of this study, only one *MAT* allele was known for each of the *Trichosporonales* species. To verify that both *MAT* alleles, A1 and A2, can be found in different strains within a single species, we analyzed additional *Vanrija humicola* strains (Table 2). *V. humicola* is a member of a phylogenetically early-branching lineage within the *Trichosporonales*, and the previously sequenced strain (JCM1457) carries the A2 allele (Figs 1 and 2) [51]. Several other strains obtained from culture collections were analyzed by RAPD (Rapid Amplification of Polymorphic DNA) genotyping, which revealed banding patterns similar to those from the type strain *V. humicola* JCM1457 (=CBS571, *MAT* A2) (S1 Fig). These strains were analyzed further by PCR for the mating type genes, and based on the analysis of PCR amplicons of the *HD* gene-containing regions, strains CBS4282 and CBS4283 were preliminarily identified as *MAT* A1 strains. To assess this further, we sequenced the genome of CBS4282 using Illumina sequencing. The genome was assembled into 21 scaffolds with 5089 predicted genes. At 22.63 Mb, the assembly size is similar to that of strain JCM1457 (A2) (22.65 Mb, [51]). A k-mer analysis of the Illumina reads for CBS4282 showed a single peak as expected for a haploid genome (S1 Fig). Analysis of the *MAT* region revealed fused *MAT* loci within a ∼50 kb region of scaffold 02 carrying the *SXI2* gene and *STE3*α allele, confirming that this strain is indeed an A1 strain, and thus showing that alternative alleles can be found in different strains of a single *Trichosporonales* species (Figs 2 and 3, S2 and S3 Figs, Table 2). This was further confirmed by an analysis of the recently published genome of a second *Apiotrichum porosum* strain [52], with the two *A. porosum* genome sequences available now representing the A1 and A2 alleles (Figs 2 and 3).

**Table 2.** Strains used in this study.

| **Species** | **Strain** | ***MAT* allele** | **Source** |
| --- | --- | --- | --- |
| *Cutaneotrichosporon oleaginosum* | ATCC20508 | A2 | American Type Culture Collection |
| *Vanrija humicola* | JCM1457 (= CBS571) | A2 | Westerdijk Fungal Biodiversity Institute |
| *Vanrija humicola* | CBS4281 | A2 | Westerdijk Fungal Biodiversity Institute |
| *Vanrija humicola* | CBS4282 | A1 | Westerdijk Fungal Biodiversity Institute |
| *Vanrija humicola* | CBS4283 | A1 | Westerdijk Fungal Biodiversity Institute |
| *Vanrija humicola* | CBS8354 | A2 | Westerdijk Fungal Biodiversity Institute |
| *Vanrija humicola* | CBS8371 | A2 | Westerdijk Fungal Biodiversity Institute |

**Fig. 3.**
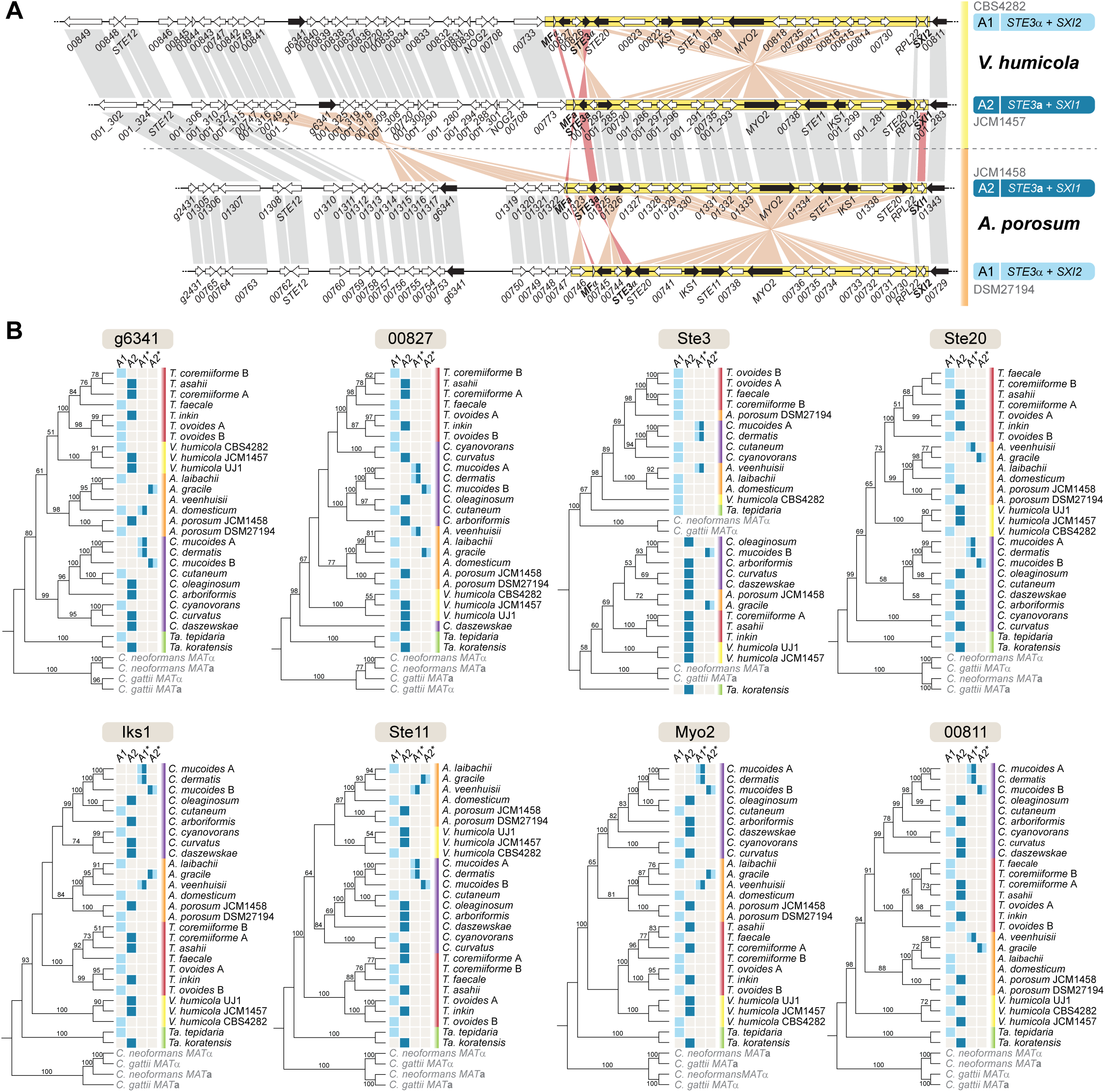
Phylogenetic analysis of proteins encoded by genes within and around the *MAT* loci in *Trichosporonales*. **A.** The A1 and A2 *MAT* alleles of *V. humicola* and *A. porosum* are shown for reference of the positions of the analyzed genes within the *MAT* region (depicted by black arrows). **B.** Individual genealogies of the selected genes across species. In the core *MAT* region (enclosed in yellow in panel A), a deep trans-specific polymorphism is only seen for the *STE3* gene. In *V. humicola*, most of the analyzed genes within this region branch by mating type. The *MAT* alleles of the analyzed strains are given next to the phylogeny. Genus name abbreviations and other color features are given as in Figures 1 and 2.

### Alternative *MAT* alleles in *Trichosporonales* show significant chromosomal rearrangements, but each displays highly conserved structure across species

The gene order within the core *MAT* region between the *SXI2* and *STE3*α/*MF*α genes is very well conserved between *V. humicola* CBS4282 (A1) and the A1 alleles of the other *Trichosporonales* species analyzed, whereas it differs from the *V. humicola* A2 strain JCM1457 by two inversions, one of which encompasses most of the core *MAT* region (Fig 2). The same is true for the A2 alleles, where the core *MAT* region is also rather conserved between *V. humicola* JCM1457 (A2) and other *Trichosporonales* (Fig 2).

In addition to the key mating-type determinants, the *MAT* loci of the *Trichosporonales* contain other genes previously shown to be required for mating and filamentation in other species (e.g. *STE11, STE12* and *STE20*), and their common presence in the *MAT* loci of *Tremellales*, suggests these genes were anciently recruited to the *MAT* locus (Fig 2). In the *Trichosporonales, STE11* and *STE20* are found within the core *MAT* region (between the *HD* and *P/R* genes), and *STE12* is in the vicinity of the core *MAT* region in most species (Fig 2). One exception turned out to be *STE12* in the *Cutaneotrichosporon* lineage. In *C. oleaginosum* strains IBC0246 (A2) and ATCC20509 (A2), *STE12* is located on a different scaffold than the *MAT* locus. While *STE12* in IBC0246 (A2) is located at the end of its respective scaffold, its location in ATCC20509 (A2) is around position 1.1 Mb of a 2.5 Mb scaffold, indicating that even if this scaffold was linked to the *MAT* scaffold, the linkage would not be very tight. To confirm that *STE12* is indeed not linked to the *MAT* locus in *C. oleaginosum*, we sequenced the genome of the third of the three known strains of this species, ATCC20508. Analyses of the ATCC20508 (A2) and ATCC20509 (A2) genome assemblies showed that both the *MAT* locus and *STE12* are located within large co-linear scaffolds in both strains, providing further evidence that *STE12* is not linked to the *MAT* locus in this species (Fig S4). Consistent with this, in the other *Cutaneotrichosporon* species analyzed, *STE12* is either on the same scaffold as *MAT*, but at a distance of 400 to 750 kb (S5 Fig), or is present on different scaffolds than *MAT*. Thus, while this suggests that the location of *STE12* in the vicinity of *MAT* likely represents the ancestral state in the *Trichosporonales*, its relocation to a different genomic location in the *Cutaneotrichosporon* species indicates that *STE12* might not be fully linked to *MAT* in the *Trichosporonales*.

Another difference in *MAT* organization that can be observed in the genus *Cutaneotrichosporon* is the combination of *HD* genes and *STE3* alleles. While all other strains of each species carry either the A1 (*STE3*α+*SXI2*) or A2 (*STE3***a**+*SXI1*) *MAT* loci, *Cutaneotrichosporon dermatis* carries an allele that has the overall genomic organization of A1, but has instead a *SXI1* gene (S5 Fig). Because the A1 vs. A2 designation is based on the *STE3* variant, this allele was called A1*. Interestingly, A1* alleles can also be found in *Apiotrichum veenhuisii*, whereas a corresponding A2* allele, in which *STE3***a** is linked to the *SXI2* gene, can be found in *Apiotrichum gracile* (Fig 1 and S5 Fig). Furthermore, both A1* and A2* alleles can be found in *Cutaneotrichosporon mucoides*, one of several hybrid species within the *Trichosporonales* (S6 Fig, S2 Text) [50, 51]. These *MAT* alleles seem to represent a derived state and might have been independently generated by recombination involving crossing-over between the *MAT* region adjacent to the *SXI1*/*SXI2* genes that is co-linear between the A1 and A2 alleles (S5 Fig).

### Potential ancient origin of the fused *MAT* loci in *Trichosporonales*

The nearly identical gene content and overall high degree of conservation in gene order and allele combinations of *HD* and *P/R* loci in the *Trichosporonales* suggests that the fusion of the *MAT* loci most likely occurred in the common ancestor of the *Trichosporonales* lineage, although other hypotheses are possible, which are discussed below. To determine at what time this fusion might have occurred, we estimated divergence times of the basidiomycete lineages included in Figure 1 using three calibration points (see S7 Fig and Materials and Methods). Based on this analysis, the *Trichosporonales* and *Tremellales* have a common ancestor dating back approximately 179 million years ago (MYA), whereas the earliest split within the *Trichosporonales* occurred approximately 147 MYA (S7 Fig). Thus, under the assumption of a *MAT* fusion in the common ancestor of the *Trichosporonales*, this fusion would have occurred between 179 and 147 MYA (S7 Fig). While our estimates of divergence times for the *Microbotryum* and *Trichosporonales* lineages agree well with recent analyses [8, 54], it should be noted that previous estimates for the last common ancestor of the pathogenic Cryptococci resulted in earlier divergence times ranging from 40 to 100 MYA [55-58]. Nevertheless, even if the divergence time of the ancestor of the pathogenic Cryptococci was underestimated in our analysis, this would still place the last common ancestor of the *Trichosporonales*, and thus the likely *MAT* fusion in this group, at a much earlier time point.

### Analysis of recombination suppression in the fused *MAT* loci in *V. humicola*

Suppression of recombination is a hallmark of sex chromosomes of animals and plants and the mating-type chromosomes of algae and fungi [6, 10, 26, 32, 35, 59, 60]. In basidiomycetes, recombination cessation between *P/R* and *HD* loci is often established following fusion (i.e. the physical linkage) of the two loci, and sometimes expanding far beyond the physically linked *MAT* loci [8, 26]. One consequence of recombination cessation can be the accumulation of transposable elements as well as increased genetic differentiation between allelic sequences [27]. This was observed in *C. neoformans*, where the *MAT***a** and *MAT*α alleles differ significantly in gene organization, and the *MAT* locus contains more remnants of transposons and other repeat sequences than other genomic regions except for the centromeres and rDNA repeats [24, 61]. Furthermore, both *C. neoformans MAT* alleles are highly rearranged in comparison with their *Cryptococcus gattii* counterparts [35].

In contrast, the *MAT* A1 and A2 alleles of *V. humicola* are overall co-linear apart from two inversions, and this is generally the case throughout the *Trichosporonales* (Figs 2 and 3A). An analysis of repetitive sequences shows that there is no accumulation of repeat regions within the *MAT* alleles of *V. humicola* (S1 Table). Thus, while the two inversions might impair meiotic pairing and therefore recombination in this region, the conserved gene order and lack of repeats made us consider if the *V. humicola MAT* loci, and more generally the *Trichosporonales MAT* loci, are regions of suppressed recombination.

To test this, we first analyzed phylogenetic trees for several genes present within or adjacent to the *MAT* loci of *Trichosporonales* (Fig 4). For genes in regions undergoing meiotic recombination, alleles associated with alternative mating types are expected to display a species-specific topology in a phylogenetic tree, whereas genes in *MAT* regions should cluster by mating type if recombination suppression predates speciation (i.e. with the A1 alleles of the different species branching together rather than each of the alleles clustering with the A2 allele from the same species) [31, 35]. In the *Trichosporonales*, only *STE3* clearly shows a mating type-specific pattern with an ancient trans-species polymorphism in *Trichosporonales* and *Tremellales* (Fig 3B). None of the other genes tested showed a mating type-specific phylogenetic pattern for the *Trichosporonales* at such a deep phylogenetic level, indicating that recombination was not suppressed at the base of the clade. However, for the genes *STE20, IKS1, STE11*, and *MYO2* within the core *MAT* locus, sequences from two *V. humicola* A2 strains (JCM1457 and UJ1) group separately from the A1 sequence (CBS4282) (Fig 3B). Therefore, these genes might have evolved mating type-specific alleles at the species level, although more strains from more species would have to be investigated to exclude this finding occurring by chance (Fig 3).

**Fig 4.**
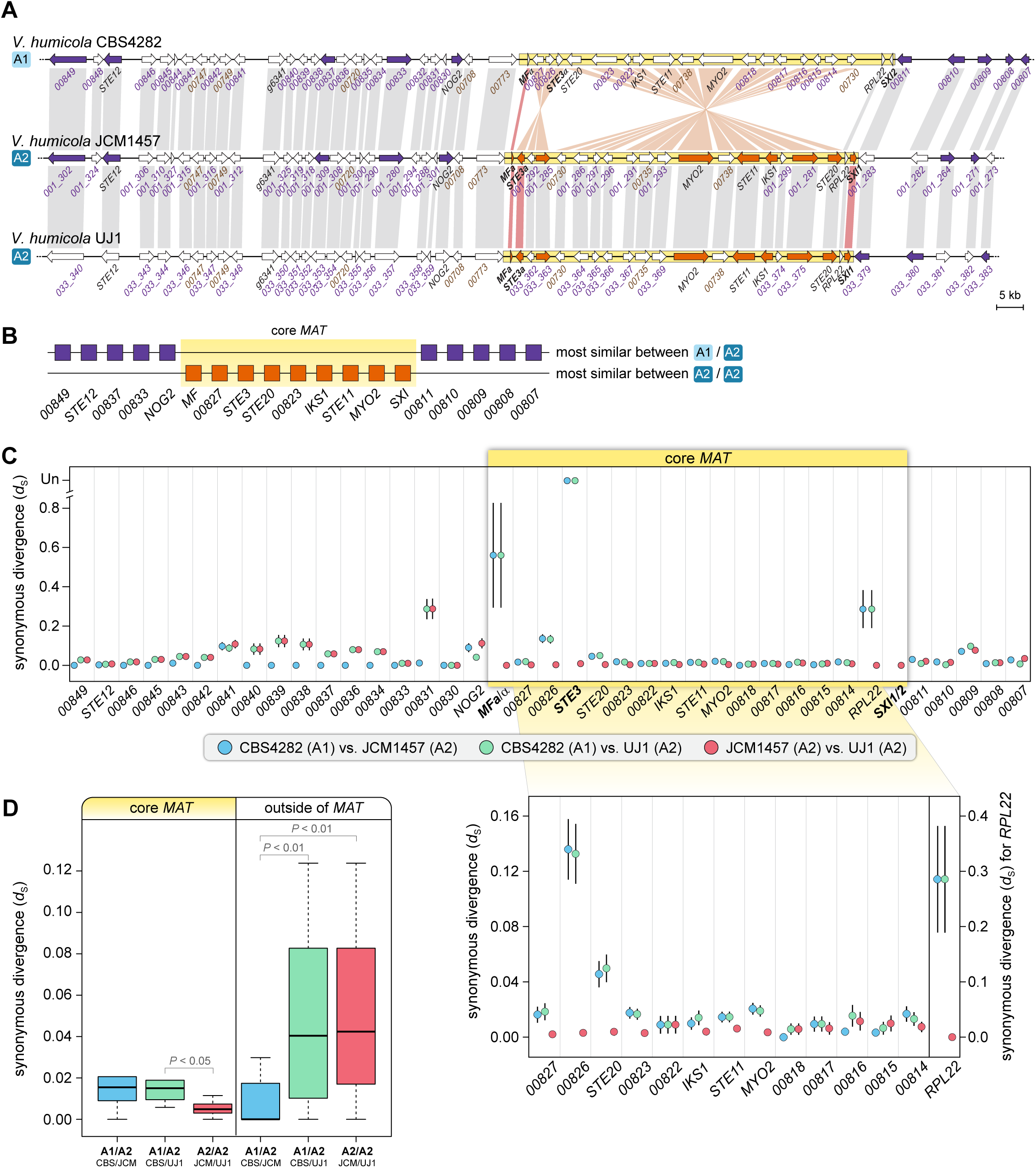
*MAT* alleles of *V. humicola* strains. **A.** Two strains (JCM1457 and UJ1) are *MAT* A2, while strain CBS4282 is a *MAT* A1 strain. Genes shown in the same color (orange within the core *MAT* region, and purple outside of the *MAT* region) were the most similar pair in a pairwise BLASTN analyses among the three alleles of the different *V. humicola* strains. **B.** Schematic view of BLASTN results. For each analyzed gene, it is shown if alleles of the most similar pair came from the strains with the same (A2/A2) or different (A1/A2) *MAT* alleles. Within the core *MAT* region, higher sequence similarity was observed between the two A2 alleles than between A1 and A2 alleles. **C.** Synonymous divergence between *MAT* alleles in different *V. humicola* strains. Synonymous substitutions per synonymous site and standard errors (*d*_S_ +/-SE) are shown for pairwise comparisons between strains CBS4282 (A1), JCM1457 (A2), and UJ1 (A2) for genes in the region containing the mating type genes. For *STE3* genes, for which no *d*_S_ value could be calculated in comparisons of strains with different mating types, values are shown as undefined (Un) at the top of the diagram. The inset below shows *d*_S_ values in the region between the core *MAT* genes, gene order as in strain CBS4282 (A1), *STE3* is left out for clarity. **D.** Boxplots of distribution of *d*_S_ values shown in C (only genes for which values could be calculated were used in this comparison). The plots show the distribution of *d*_S_ values, with median value as a horizontal line in the box between the first and third quartiles. Outliers were left out for better visibility. Significantly different values are indicated (p-values from Student‘s t-test).

The possibility of mating type-specific alleles is further supported by BLASTN analyses comparing alleles in the three available *V. humicola* genomes, which showed that within the core *MAT* region, alleles from the A2 strains (JCM1457 and UJ1) are more similar to each other than to the A1 strain CBS4282, whereas outside of this region this is not the case (Figs 4A and 4B). This suggests that the *MAT* locus of *V. humicola* might be a region of recent recombination suppression.

To further test whether recombination is suppressed between the *MAT* alleles in *V. humicola*, we analyzed levels of synonymous divergence (*d*_S_) between alleles on the *MAT*-containing scaffold as well as on three other scaffolds of CBS4282 (A1) and the two *V. humicola* A2 strains (S8 Fig). In the four longest scaffolds, genes within large regions of the scaffolds are more similar between CBS4282 (A1) and the JCM1457 (A2) than between the two A2 strains, where stretches of low *d*_S_ values occur much less frequently (S8 Fig). Interestingly, we also noted regions of high similarity between CBS4282 (A1) and JCM1457 (A2) that are interrupted by stretches containing more divergent alleles. This pattern is consistent with ongoing recombination in natural *V. humicola* populations, even though sexual reproduction has not yet been observed.

In contrast to other genomic regions, including the co-linear regions outside of the core *MAT* region where overall divergence is lowest between CBS4282 (A1) and JCM1457 (A2) (Fig 4C and S8 Fig), most of the genes in the core *MAT* region show slightly higher levels of divergence between the A1 strain and the two A2 strains than between the two A2 strains (Fig 4C). This is consistent with an absence of genetic exchange between the A1 and A2 alleles, as one would expect if recombination in this region is suppressed. These findings might be explained by a reduced recombination rate in the core *MAT* region carrying the inversions in alternate *MAT* alleles, which could lead to accumulation of mutations in the two *MAT* alleles and thus to elevated synonymous divergence between A1 and A2 alleles. However, divergence between A1 and A2 strains within the core *MAT* region is only moderately elevated for most genes, and the difference between the average *d*_S_ values is statistically significant (P< 0.047, t-test statistics 1.8101) only when comparing the analysis of CBS4282 (A1) vs. UJ1 (A2) with the analysis of JCM1457 (A2) vs. UJ1 (A2) (Fig 4D). One possible explanation could be that genetic exchange within the inverted regions may not be completely inhibited, because exchange via non-crossover gene conversion or double crossover can still occur within the inverted regions [62]. This should result in a so-called suspension bridge pattern with divergence in the middle of an inverted region lower than towards the inversion breakpoints [63]. To test this, we performed BLASTN analyses using sliding windows of genomic sequences of the *MAT* locus and adjacent regions (S9 Fig). The results show less sequence similarity in the regions of putative inversion breakpoints between A1 and A2 strains compared to regions within the inversions and outside of the *MAT* region, consistent with the hypothesis that a certain amount of recombination is occurring within the inverted regions.

The incomplete suppression of recombination (and thus the possibility of ongoing genetic exchange) might be a reason why the gene order within each A1 and A2 allele in different species is surprisingly well conserved (Fig 2). To test if this degree of gene order conservation extends beyond the *MAT* locus, we compared the *MAT*-containing scaffolds of *Trichosporonales*. As expected for a range of species separated by millions of years of evolution, synteny is conserved between closely related species, but not between species that diverged long ago (S10 Fig), except for the *MAT* region. This suggests that the *MAT* region of *Trichosporonales* is an ancient cluster of tightly linked loci (also known as ‘supergenes’) that seem to segregate as a stable polymorphism within the populations of each species. Thus, the *Trichosporonales* appear to contain very stable *MAT* alleles with respect to gene order, combined with (slightly) suppressed recombination between different alleles.

### The *MAT* loci in the *Trichosporonales* have a significantly lower GC content compared to the overall genomic GC content

One curious observation was found during the analysis of the *Trichosporonales MAT* region, namely a lower GC content in the *MAT* regions compared to the surrounding regions for the analyzed *Vanrija, Cutaneotrichosporon*, and *Trichosporon* strains (S10 and S11 Figs). One possible explanation for a lower GC content could be an accumulation of AT-rich transposable elements, but an analysis of repeats in the three *V. humicola* strains showed that there are only a few (strain CBS4282, A1) or no (strains JCM1457 and UJ1, both A2) repeats present within the *MAT* region of these strains (S1 Table). Another explanation might be a lower density of coding regions, which tend to have a higher GC content than non-coding regions. However, an analysis of the GC content only in the coding sequences within and around the *MAT* region of strain CBS4282 showed the same pattern of lower GC content within the *MAT* region (S11B Fig). Another possible explanation for the lower GC content might be the accumulation of mutations due to reduced recombination. Under the (simplistic) assumption that mutation frequencies for all nucleotide exchanges are similar, this would drive the GC content towards 50%. In genomes with an average GC content of more than 50%, this would appear as a region with lower GC content, and this would apply to the analyzed strains with an average genomic GC content of 58 to 63% (S11 Fig). It has been shown that spontaneous mutations tend to be AT-biased in many species including several fungi [64-68], and thus an increased accumulation of mutations would generally lead to a lower GC content in the corresponding regions. Both explanations can be supported by an analysis of the codon usage within the *MAT* region of the three *V. humicola* strains (S2 Table). Among the codons for amino acids that can be encoded by more than one codon, there is a trend for GC-rich codons to be used less frequently in the *MAT* region compared to the rest of the genome, consistent with AT-biased (or GC content-equalizing) mutations combined with selection for conserved protein sequences due to functional constraints. Additional studies of species with a GC content of less than 50% might be useful to test these hypotheses.

## Discussion

### Fused *MAT* loci evolved several times in basidiomycetes

In this study, we analyzed *MAT* loci from the *Trichosporonales*, and found that all of the analyzed species harbor fused *MAT* loci with a single *HD* gene, an arrangement that among the *Tremellomycetes* has so far only been found in the pathogenic Cryptococci [18]. Physically linked *MAT* loci have been identified previously in other basidiomycetes (*Malassezia, Microbotryum, Sporisorium, Ustilago*), but these *MAT* loci carry the ancestral arrangement of two *HD* genes except in *Microbotryum violaceum caroliniana*, in which the *HD1* gene was specifically lost from the A2 mating type [8, 18, 20-23, 69, 70]. Fused *MAT* loci in fungi can be considered supergenes, which are linked genetic loci that facilitate the co-segregation of beneficial allele combinations [59]. Supergenes have evolved in a number of species, and can regulate different traits, e.g. mimicry in butterflies, social organization in ants, or mating in fungi [8, 71, 72]. Our findings add to the growing number of instances of supergene formation for fungal *MAT* loci, even though the molecular mechanisms maintaining supergenes, e.g. suppressed recombination, might not be identical in all cases as discussed below.

An analysis of several strains from *V. humicola* showed that there are two main *MAT* alleles (i.e. two main allelic combinations at the fused *MAT* loci), and both alleles are distributed throughout the *Trichosporonales* phylogenetic tree. Additional strains would have to be analyzed to establish if these two alleles are the most prevalent in this group. However, if the mating system is predominantly inbreeding, rare alleles have no advantage and can be gradually lost by genetic drift. Thus, fusion of *MAT* loci, which can evolve by selection because it is beneficial in selfing mating systems, is predicted to lead to a reduction in the number of *MAT* alleles [4, 73]. In addition, theoretical modelling predicts that a combination of facultative and rare sexual reproduction, low mutation rates, and a small effective population size should also lead to a reduction in the number of *MAT* alleles [74]. Population sizes and mutation rates are, however, not known for *Trichosporonales* species, and no sexual development has been observed so far making it possible that asexual reproduction is the predominant form of propagation for many *Trichosporonales*. Another possibility would be that only two alleles per *MAT* locus existed in a tetrapolar ancestor possibly due to other factors, such as linkage of the *MAT* loci to their respective centromeres and concomitant loss of allele diversity, as recently proposed for tetrapolar *Microbotryum* species [75]. Analyses of species branching at the base of the *Trichosporonales* lineage could help to clarify this hypothesis. Another explanation for the observed prevalence of only two *MAT* alleles might be that the *HD* and *P/R* loci are located at the two ends of the *MAT* region with an inversion between opposite alleles. Therefore, an odd number of crossovers within the region between *HD* and *P/R* would result in genetically imbalanced progeny, which would be inviable, whereas an even number of crossovers would not result in allele exchange at the *HD* or *P/R* loci. Thus, recombination could occur, but would most likely not change the *HD* and *P/R* allele combinations. In the case of the A1^*^ and A2^*^ alleles, recombination might have occurred in the rather small co-linear genomic region between the *SXI1/SXI2* genes and the adjacent inversion, and thus might be a rare event.

### Convergent evolution of fused *MAT* loci in *Trichosporonales* and *Tremellales*

Overall, our finding of fused *MAT* loci with a single *HD* gene in all of the *Trichosporonales* species investigated is most consistent with the hypothesis that a fusion event of the *HD* and *P/R* loci followed by loss of one *HD* gene per *MAT* allele occurred independently in the *Trichosporonales* and the pathogenic Cryptococci lineages. The alternative hypothesis that the fusion of the *HD* and *P/R* loci occurred in the common ancestor of the *Trichosporonales* and pathogenic Cryptococci is less parsimonious as it would imply multiple independent reversions to tetrapolarity in the non-pathogenic Cryptococci. In addition, while several mating-associated genes can be found in the *MAT* loci of both groups, the majority of genes within the *MAT* locus of the *Trichosporonales* is not found in the *MAT* locus in the *Tremellales*, consistent with independent fusion events in the two lineages. This hypothesis is also supported by the combination of the *SXI1*/*SXI2* genes and the *STE3***a**/α alleles in the A1 and A2 alleles of the *Trichosporonales*, which is different from those of the *MAT***a** and *MAT*α alleles of the pathogenic Cryptococci. A model is proposed in Figure 5 to explain the current situation in the *Tremellomycetes*, where recruitment of several mating-associated genes into the *P/R* loci of a tetrapolar ancestor occurred first as these genes can also be found within the *P/R* loci of extant *Tremellales* with unlinked *MAT* loci [30-32]. After the split of the *Tremellales* and *Trichosporonales* lineages, fusion of the *HD* and *P/R* loci as well as the loss of one *HD* gene per *MAT* allele occurred in the ancestor of the *Trichosporonales*, whereas within the *Tremellales* this happened only in the ancestor of the pathogenic Cryptococci.

**Fig 5.**
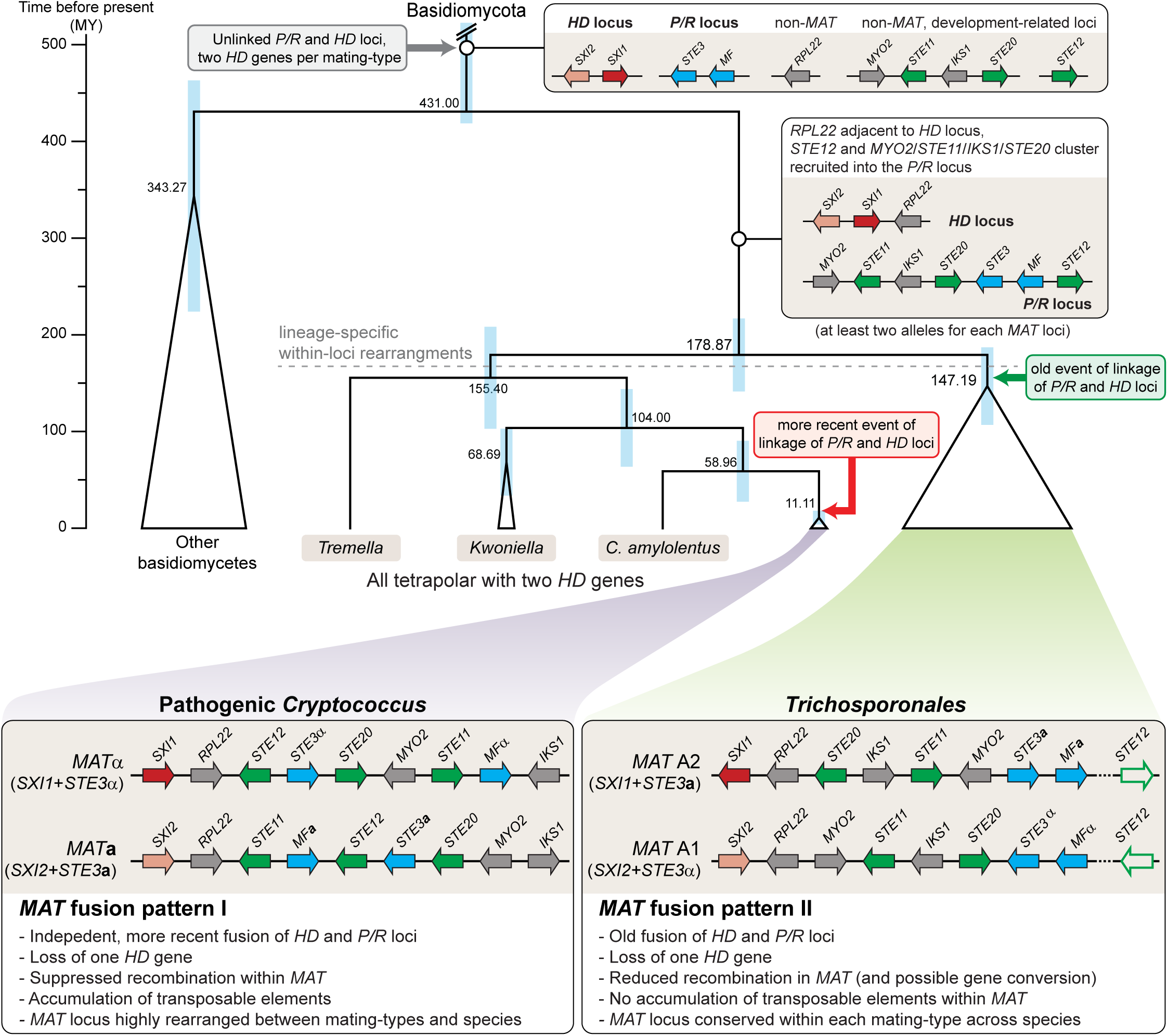
Model for the evolution of the *MAT* loci in *Tremellomycetes*. Genes at the *MAT* loci containing homeodomain transcription factor genes (*HD* locus) or pheromone and receptor genes (*P/R* locus) are shown in red/pink and blue, respectively. Genes involved in sexual development, but not originally part of a *MAT* locus are shown in green, other genes are shown in grey. Only genes from the *C. neoformans MAT* locus that are also linked to the core *MAT* genes (*STE3, HD* genes) in the *Trichosporonales* are shown (*STE11, STE12, STE20, IKS1, MYO2, RPL22*). Other genes present at the *MAT* loci are left out for clarity. A trend towards integrating other developmental genes into the *MAT* loci is reflected in the recruitment of the *STE12* gene and an ancestral *STE11*/*STE20* cluster into the *P/R* locus. This resulted in a tetrapolar arrangement with (at least) two alleles for each *MAT* locus. In the *Tremellales*, tetrapolarity is retained in the extant *Tremella, Kwoniella*, and non-pathogenic *Cryptococcus* lineages, whereas in the pathogenic Cryptococci, the *MAT* loci fused and one *HD* gene was lost from each *MAT* allele. Fusion of the *MAT* loci occurred independently and earlier in the ancestor of the *Trichosporonales*, also with loss of one *HD* gene, resulting in two known *MAT* alleles with combinations of *SXI1*/*SXI2* and *STE3***a**/*STE3*α that differ from those in the known *Cryptococcus* alleles. The *STE12* gene is shown in outline only to indicate that it is not part of the *MAT* locus in all species; *STE12* may have been recruited to the *P/R* locus in the *Tremellales-Trichosporonales* common ancestor and eventually evicted from the *MAT* locus in the *Trichosporonales* possibly associated with the fusion event of *P/R* and *HD* loci, or instead *STE12* was recruited to *MAT* only in the *Tremellales*. The phylogenetic relationships and inferred dates of speciation (numbers in black on tree nodes) are depicted according to S7 Fig. The blue bars correspond to 95 % confidence intervals.

An independent fusion might suggest that similar selective pressures were acting to result in similar evolutionary trajectories in the *Trichosporonales* and pathogenic Cryptococci. In the latter group, it has been hypothesized that the pathogenic lifestyle might make it difficult to find a mating partner while associated with a host. In this case, linkage between the two *MAT* loci is expected to be favored by selection, as this increases the odds of compatibility between the gametes derived from a single diploid zygote [4, 8, 9, 18, 26]. Extant members of the *Trichosporonales* can be associated with hosts as commensals or pathogens, but this lineage also includes many soil- or water-associated saprobes [37, 38]. It is possible that the ancestor in which *MAT* fusion occurred was associated with a host, and that the physically linked *MAT* loci remained stable during evolution of species with saprobic lifestyles. However, a scarcity of compatible mating partners might also occur under other conditions, e.g. if a population is derived from few progenitor cells that propagated through mitotic cell divisions during favorable conditions and switched to sexual development when nutrients were depleted.

An alternative hypothesis to the fusion of the *MAT* loci occurring in the common ancestor of the *Trichosporonales* would be multiple independent fusions in different lineages within the *Trichosporonales*. However, in this case, one would expect different genomic arrangements, similar to what has been observed in several cases of convergent evolution of fused *MAT* loci in the genus *Microbotryum* [8]. It is, in principle, possible that there is strong selective pressure that favors the specific genomic configurations observed in the *Trichosporonales MAT* loci for the A1 and A2 alleles, but the nature of this selective pressure, which would have to act on multiple species in different environments, is not clear. Another hypothesis could be that the fusion of *MAT* loci is ancient, but the inversion, which may have contributed to limit or repress recombination, occurred recently and independently in several *Trichosporonales* lineages. This would explain the lack of trans-species polymorphisms observed for most of the genes at the fused *MAT* loci. Such a scenario of multiple independent inversions would also require strong selective pressure in multiple species to yield not only functionally similar, but essentially the same genomic configuration. Under the hypotheses of more recent, independent fusions or inversions, one might expect to find species within the *Trichosporonales* lineage that have not undergone *MAT* fusion (i.e. physical linkage) or inversions, similar to the genus *Microbotryum*, where tetrapolar and bipolar species are intermingled in the phylogenetic tree [8]. So far, such species have not been identified in the *Trichosporonales*. Therefore, the hypothesis of a fusion event in the last common ancestor of the *Trichosporonales* appears to be the most parsimonious at present, despite several unusual genomic features discussed below.

In addition to the fused *MAT* loci, the presence of only one *HD* gene per *MAT* allele in all examined *Trichosporonales* is similar to pathogenic Cryptococci. This was noted in the initial analysis of genome sequences for two *Trichosporonales* strains now designated as *MAT* A2 [44], and the current study shows that this is a conserved feature throughout the *Trichosporonales*. Outside of the *Tremellomycetes*, the majority of analyzed basidiomycete species harbor at least two *HD* genes per *MAT* allele. In the case of two *HD* genes, one belongs to the *HD1* class, and the other to the *HD2* class, whereas in the *Agaricomycetes*, species with more than one *HD1* or *HD2* gene per *MAT* allele are known [18]. The *HD1* and *HD2* genes from the same *MAT* allele are not compatible with each other, irrespective of whether the mating system is tetrapolar or bipolar [16, 18]. An exception is the genus *Wallemia*, where fused *MAT* loci exist with only one *HD* gene, but until now, only one *MAT* allele harboring a *SXI1* homolog is known in this genus, whereas another mating type appears to not contain any *SXI* gene [18, 76, 77]. It might be that mating in *Wallemia* species does not rely on *HD* compatibility. Functionally, two compatible *HD1* and *HD2* genes from different *MAT* alleles are necessary and sufficient for sexual development not only in *C. neoformans*, where only one *HD1*/*HD2* gene pair is present after mating [78, 79], but also in *U. maydis* and *Coprinopsis cinerea*, where two (*U. maydis*) or more (*C. cinerea*) compatible *HD1*/*HD2* combinations can be present after mating [80-82]. The presence of multiple *HD* gene paralogs or increased *HD* allele diversity within a species is advantageous under outcrossing, because it allows more frequent dikaryon viability after mating when the *P/R* locus also exists in multiple alleles [16]. However, if a species is predominantly selfing, a single *MAT* locus with two alleles will provide the highest percentage of compatibility between gametes from a single tetrad [4, 16, 18]. In such cases, the loss of one *HD* gene per *MAT* locus should not be a problem from either a functional or evolutionary point of view, and thus this genomic configuration might be observed in other species with fused *MAT* loci, unless the *HD* genes have other functions unrelated to mating. Another possible reason for the scarcity of *HD* loci with only a single *HD* gene might be that having two different *HD* genes occupying allelic positions results in a genomic region that might not be able to properly pair during meiosis. In the ascomycete *Neurospora crassa*, unpaired regions are known to trigger MSUD (meiotic silencing of unpaired DNA) through an RNAi-based mechanism, although interestingly the mating type genes of *N. crassa* are immune to silencing in their normal genomic location [83-85]. Thus, it might be possible that *MAT* loci with a single *HD* gene have properties at the DNA level that are independent of the functions of the encoded proteins.

One common feature of fungal *MAT* loci and sex chromosomes in other eukaryotes might be the recruitment of sex-associated genes. The presence of the *STE11* and *STE20* within the core *MAT* loci of *Trichosporonales* and *Tremellales* suggests an ancient linkage of these genes to the *MAT* locus. In *C. neoformans, STE20* is required for the formation of proper heterokaryotic filaments and basidia after mating, and *STE11*α is required for mating and filament formation [86, 87]. *STE20* displays *MAT*-specific alleles [30, 31] and is also located within the *P/R* locus of other basidiomycete species, e.g. *Leucosporidium scottii* and red yeasts in the *Pucciniomycotina* [17]. However, the functions of many genes within the *MAT* loci are not known, and further experiments will be needed to determine if these genes have functions in mating or sexual reproduction.

### Genomic signatures that distinguish the *MAT* loci from surrounding genomic regions in *Trichosporonales*

A common feature of fungal *MAT* loci and the sex-determining regions in other eukaryotic groups is that recombination is usually suppressed in these regions. In several fungi, suppressed recombination is observed not only within the *MAT* locus itself, but spreading out from the *MAT* locus along the *MAT*-carrying chromosome resulting in so-called evolutionary strata of stepwise recombination suppression, similar to findings in sex-chromosomes of mammals and plants. Examples are the ascomycete *Neurospora tetrasperma* and the basidiomycete genus *Microbotryum* [8, 10, 26, 29, 88-92]. In the latter, it was recently shown that the linkage of *HD* and *P/R* loci evolved independently at least five times within the genus, followed in each case by further stepwise expansions of the regions of suppressed recombination beyond the fused *MAT* loci, possibly through neutral rearrangements [8, 26]. In contrast, the analysis of *V. humicola* did not yield indications for suppressed recombination beyond the core *MAT* region that lies between the *SXI1*/*SXI2* and pheromone genes. Furthermore, the gene content and gene order within the *MAT* loci and the *HD*/*STE3* allele combinations in the analyzed *Trichosporonales* species are largely conserved, suggesting that the fusion of *HD* and *P/R* loci occurred once in a common ancestor. Regions outside of the *MAT* locus have generally undergone substantial rearrangements (e.g. translocations) in different species as seen by the varying genomic locations of *STE12* in the genus *Cutaneotrichosporon* and the lack of synteny outside of the *MAT* region in more distantly related *Trichosporonales*.

In *Microbotryum*, it was shown that evolutionary strata ranging from mating-type- to species-specific patterns can be distinguished along chromosomes harboring recently linked *MAT* loci [8, 26]. These analyses were performed based on alleles from several strains per species and of both mating types. One challenge for the phylogenetic analysis in the *Trichosporonales* is that most species are currently represented by a single strain. Analyzing additional alleles from more strains of each species will be necessary to fully evaluate the phylogeny of the genes located within and surrounding the *MAT* locus.

In the *Trichosporonales*, the two *MAT* alleles, A1 and A2, differ in general by two inversions. If the *MAT* fusion is relatively ancient, i.e. occurred in the last common ancestor of the *Trichosporonales*, a remaining question is why the genetic differences in the two known alleles are not more pronounced. One explanation could be that recombination within the inverted regions might still be possible via non-crossover gene conversion or double crossover [63]. This would be consistent with the pattern observed in *V. humicola* of lower divergence between A1 and A2 alleles within the inverted regions compared to divergence at the inversion breakpoints. In *C. neoformans*, gene conversion occurred in a GC-rich intergenic region within the *MAT* locus and was proposed as a mechanism for maintaining functionality of those genes within the *MAT* locus that are essential [36]. Gene conversion has also been observed in regions with suppressed recombination in the mating-type chromosomes of *N. tetrasperma*, in the mating type locus of the green algae *Chlamydomonas reinhardtii*, and in sex chromosomes of animals [93-99].

Gene conversion might be associated with an increase in GC content as it has been observed in a number of eukaryotes that gene conversion can be biased towards GC [100-102], which would not be compatible with the observed lower GC content in the *MAT* region of *Trichosporonales*. However, a recent analysis of two fungal species found no evidence of a GC-bias in gene conversion [103], and therefore at present gene conversion as an explanation for low divergence between *MAT* alleles in *Trichosporonales* remains possible. It is tempting to speculate that the lower GC content observed in the *MAT* region of *Trichosporonales* is connected to the incomplete suppression of recombination and/or the (yet unknown) mechanisms that lead to the low levels of divergence between the *MAT* alleles. In *C. neoformans*, two regions of higher GC content outside of the *MAT* locus are associated with increased recombination [104]. A correlation between higher GC content and increased recombination has been demonstrated in humans, although it is not clear in this case what is cause and what is effect, whereas in the yeast *Saccharomyces cerevisiae*, a correlation was described at the kilobase range, but no long-range correlation was found [105-107].

Another hypothesis to explain the low levels of divergence between the *MAT* alleles could be same mating-type mating (“unisexual reproduction”, i.e. sexual reproduction without the need of a partner with a different mating type), which occurs in several pathogenic Cryptococci and the human pathogenic ascomycete yeast *Candida albicans* [108, 109]. Similar to reproduction after fusion of *MAT***a** and *MAT*α cells, “unisexual reproduction” in Cryptococci also entails diploidization and meiosis including genetic exchange [109-111]. Diploidization can occur without cell fusion through endoreplication, or through fusion of two cells carrying the same *MAT* allele [112]. In the latter case, this would allow recombination within the *MAT* region, which could prevent degeneration of the *MAT* locus despite suppressed recombination when paired with a different *MAT* allele. Recombination within the *MAT* locus during “unisexual reproduction” was shown for *C. neoformans* [113]. So far, no sexual cycle has been observed in the *Trichosporonales*, and therefore it is not known what forms of sexual reproduction occur in this group. This is an important open question that requires detailed investigation in future studies.

## Conclusions

In summary, we have shown that all analyzed *Trichosporonales* species contain fused mating type loci with a single *HD* gene per mating type. The two known *MAT* alleles differ by two inversions, but both alleles have a relatively stable gene content across different species, making it possible that the fusion of the *MAT* loci occurred in the last common ancestor of the *Trichosporonales*. An alternative hypothesis would be several independent fusion or inversion events, which could explain the lack of trans-species polymorphisms for most of the genes at the fused *MAT* loci. However, in contrast to *Microbotryum* species, for which multiple fusion events were demonstrated [8], the *MAT* loci in the *Trichosporonales* show allelic gene orders that are highly conserved across different species. While it is possible that strong selection favors only these genomic configurations, which could therefore have occurred multiple times, the more parsimonious hypothesis seems to be an ancient fusion event in the common ancestor of *Trichosporonales*. The apparent evolutionary stability of the alleles A1 and A2 through many speciation events (even though neighboring regions underwent significant recombination) suggests the presence of strong selective pressure that maintains the integrity of these alleles, probably due to retaining the advantages of physically linked *MAT* loci with few alleles. Another possible explanation might be that the observed combinations (*STE3*α and *SXI2* in the A1, and *STE3***a** and *SXI1* in the A2 allele) have a higher fitness than the alternative combinations. However, the presence of the A1* and A2* alleles in several *Trichosporonales* species shows that these combinations also occur, making this hypothesis less likely. A third possibility might be that the inversion that distinguishes A1 and A2 prevents the occurrence of the single (or other odd number) crossover required for allele exchange at the *HD* and *P/R* loci. Mechanistically, a combination of 1) gene conversion during meiosis after fusion of cells with different mating types to account for the limited suppression of recombination, and 2) meiotic recombination by crossing over during “unisexual reproduction” might explain the apparent evolutionary stability of the *MAT* region while maintaining (at least) two distinct alleles.

Based on the phylogenetic distribution, gene content, and allele combinations in fused *MAT* loci in pathogenic Cryptococci of the sister order *Tremellales*, fusion of *MAT* loci as well as the loss of one *HD* gene per *MAT* locus likely occurred independently in the *Trichosporonales* and *Tremellales*. Thus, our data support a model of convergent evolution of the *MAT* locus at the molecular level in two different *Tremellomycetes* orders, similar to patterns observed in other fungal groups [1, 8, 22, 23]. Future analyses of additional *Tremellomycetes* groups that have not yet been analyzed with respect to their *MAT* loci will determine if this proposed evolutionary route has been repeated independently in other cases. It will be especially interesting to analyze earlier-branching sister lineages to the *Tremellales* and *Trichosporonales* with respect to their *MAT* loci configurations and the number of *HD* genes per *MAT* locus. Other important questions are the frequency and mechanisms of recombination in the region between the fused *HD* and *P/R* loci. These questions can be addressed once *Trichosporonales* species have been identified that undergo sexual reproduction in the laboratory, so that the progeny of genetic crosses can be analyzed.

## Materials and Methods

### Strains and growth conditions

*C. oleaginosum* and *V. humicola* strains used in this study are given in Table 2. Strains were grown on YPD medium at 25°C on solid medium or in liquid culture with shaking (250 rpm) for preparation of nucleic acids.

### CHEF (contour-clamped homogeneous electric field) electrophoresis analysis of the *C. oleaginosum* isolates

CHEF plugs of the *C. oleaginosum* isolates were prepared and the electrophoresis was carried out as described in previous studies with slight modification [30, 32]. Specifically, the plugs were run using two different sets of switching time to get better resolution of the larger (S4A Fig top, 20 – 30 minutes linear ramp switching time) and smaller (S4A Fig bottom, 120 – 360 seconds linear ramp switching time) chromosomes, respectively.

### Genotyping and Analysis of mating type genes in *V. humicola* strains

A collection of *V. humicola* isolates were genetically screened by modified RAPD using primer 5’-CGTGCAAGGGAGCACC-3’ with 48°C as annealing temperature (S1A Fig). The *V. humicola* strains showing identical genotypic profile as the type strain CBS571 (Table 2) were further analyzed for the presence of the *SXI1, SXI2, STE3*, and *MYO2* genes using PCR with oligonucleotides given in S3 Table. PCR fragments were either sequenced directly or cloned into pDrive (Qiagen, Hilden, Germany) and sequenced.

### Genome sequencing, assembly, and annotation of *V. humicola* CBS4282 and *C. oleaginosum* ATCC20508

Genomic DNA samples were extracted using a modified CTAB protocol as previously reported [32, 114]. Specifically, to enrich samples with high molecular weight, after precipitation, the genomic DNA was picked out from the solution instead of spun down, and the samples were checked by CHEF for their sizes and integrity, following manufacturer’s protocol (BioRad, Hercules, CA, USA). Sequencing of the ATCC20508 and CBS4282 genomes was carried out at the Sequencing and Genomic Technologies Core Facility of the Duke Center for Genomic and Computational Biology, using large insert library (15-20 kb) and PacBio Sequel (2.0 chemistry) for ATCC20508, and generating 151 nt paired-end Illumina reads for CBS4282. The PacBio sequence reads for *C. oleaginosum* ATCC20508 were assembled using the HGAP4 assembly pipeline based on the Falcon assembler [115] included in the SMRT Link v5.0.1 software by Pacific Biosciences. The Illumina sequence reads of *V. humicola* CBS4282 were trimmed using Trimmomatic (v0.36) [116] with the following parameters to remove adapter contaminations: ILLUMINACLIP:TruSeq3-PE-2.fa:2:30:10:1:TRUE LEADING:3 TRAILING:3 SLIDINGWINDOW:4:15 MINLEN:40. Trimmed reads were assembled with SPAdes (v3.11.1) [117], and contig sequences were improved using pilon [118] based on the Illumina reads mapped to the assembly using Bowtie2 (v2.2.6) [119]. k-mer frequencies were analyzed based on the CBS4282 Illumina reads as described previously [120, 121]. Gene models for the newly sequenced strains as well as for strains for which genome assemblies, but not annotation was available from GenBank were predicted *ab initio* using MAKER (v2.31.18) [122] with predicted proteins from *C. oleaginosum* as input [44].

### Analysis of mating type regions, synonymous divergence, and repeat content

*MAT* regions in *Trichosporonales* genomes were identified by BLAST searches [123] against the well-annotated *MAT*-derived proteins from *C. neoformans* [35], and manually reannotated if necessary. The short and not well conserved pheromone precursor genes were not among the predicted genes, and were identified within the genome assemblies using custom-made Perl scripts searching for the consensus sequence M-X(15-60)-C-[ILMVST]-[ILMVST]-X-Stop.

Synteny analysis of the genomes of *V. humicola* strains and *C. oleaginosum* strains was done with nucmer from the MUMmer package (v3.23) [124]. Synteny plots were drawn with Circos [125]. Synteny between *MAT* regions of different *Trichosporonales* species was based on bidirectional BLAST analyses of the corresponding predicted proteins. For the *d*_S_ plots, alleles in *V. humicola* strains were identified by two-directional BLAST analysis, and MUSCLE (v3.8.31) [126] was used to align the two alleles per gene per strain pair. Synonymous divergence and standard errors were estimated with the yn00 program of the PAML package (v4.9) [127]. Analysis of transposable elements and other repeats in *V. humicola* genomes was performed with RepeatMasker (Smit AFA, Hubley R, Green P. RepeatMasker Open-4.0. 2013-2015, http://www.repeatmasker.org) based on the RepbaseUpdate library [128] and a library of *de novo*-identified repeat consensus sequences for each strain that was generated by RepeatModeler (Smit AFA, Hubley R. RepeatModeler Open-1.0. 2008-2015, http://www.repeatmasker.org/RepeatModeler/) as described [121].

Linear synteny comparison along the *MAT*-containing scaffolds (S10 Fig) was generated with Easyfig [129] using a minimum length of 500 bp for BLASTN hits to be drawn.

### Species tree and gene genealogies

Of the 24 *Trichosporonales* species for which genome assemblies were available we selected only 21 for phylogenetic analysis as the remaining three species (*Trichosporon coremiiforme, Trichosporon ovoides* and *Cutaneotrichosporon mucoides*) were shown to be hybrids in previous studies [50, 51]. Additionally, we selected other well-studied bipolar and tetrapolar representatives belonging to the three major Basidiomycota lineages, plus two ascomycetes as outgroup. To reconstruct the phylogenetic relationships among the selected members, the translated gene models of each species were clustered by a combination of the bidirectional best-hit (BDBH), COGtriangles (v2.1), and OrthoMCL (v1.4) algorithms implemented in the GET_HOMOLOGUES software package [130] to construct homologous gene families. The *Cryptococcus neoformans* H99 protein set was used as reference and clusters containing inparalogs (i.e. recent paralogs defined as sequences with best hits in its own genome) were excluded. A consensus set of 32 protein sequences was computed out of the intersection of the orthologous gene families obtained by the three clustering algorithms. Protein sequences were individually aligned with MAFFT v7.310 [131] using the L-INS-i strategy and poorly aligned regions were trimmed with TrimAl (-gappyout). The resulting alignments were concatenated to obtain a final supermatrix consisting of a total of 21,690 amino acid sites. We inferred a maximum-likelihood phylogeny using the LG+F+R5 model of amino acid substitution in IQ-TREE v1.6.5 [132]. Branch support values were obtained from 10,000 replicates of both ultrafast bootstrap approximation (UFBoot) [133] and the nonparametric variant of the approximate likelihood ratio test (SH-aLRT) [134].

For phylogenetic analysis of selected genes within and outside the *MAT* region in *Trichosporonales*, protein alignments were generated and trimmed as above and subsequently used to infer maximum likelihood phylogenies in IQ-TREE. Consensus trees were graphically visualized with iTOL v4.3.3 [135].

### Divergence time analysis

Divergence times were estimated in MEGA-X [136] using the RelTime approach [137]. Contrary to other methods, RelTime does not require assuming a specific model for lineage rate variation and was shown to be as accurate as other approaches using relaxed and strict molecular clock models [138]. The reconstructed species tree, with branch lengths in the units of number of substitutions per site, was used as input and transformed into an ultrametric tree with relative times. The final timetree was obtained by converting the relative node ages into absolute dates by using three calibration constraints: 0.42 million year (MY) corresponding to the divergence between *Microbotryum lychnidis-dioicae* and *Microbotryum silenes-dioicae* [139]; 41 MY for the *Ustilago – Sporisorium* split; and 413 MY representing the minimum age of Basidiomycota. The latter two calibration points were obtained from the Timetree website (http://www.timetree.org/), which should be referred to for additional information and references.

## Data availability statement

The *V. humicola* CBS4282 genome sequence (BioProject PRJNA475686) has been deposited at DDBJ/ENA/GenBank under the accession QKWK00000000. The version described here is version QKWK01000000. Illumina reads have been deposited in the NCBI SRA database under accession SRP150316. The *C. oleaginosum* ATCC20508 genome sequence (BioProject PRJNA475739) has been deposited at DDBJ/ENA/GenBank under the accession QKWL00000000. The version described here is version QKWL01000000. Pacific Biosciences reads have been deposited in the NCBI SRA database under accession SRP150334.

## Acknowledgements

The authors thank Shelby Priest for critical reading of the manuscript. MN would like to thank Swenja Ellßel and Silke Nimtz for excellent technical assistance, and Ulrich Kück and Christopher Grefen for support at the Botany Department of the Ruhr-University Bochum.

