## Supplemental figures and texts for "Convergent evolution of linked mating-type loci in basidiomycete fungi"

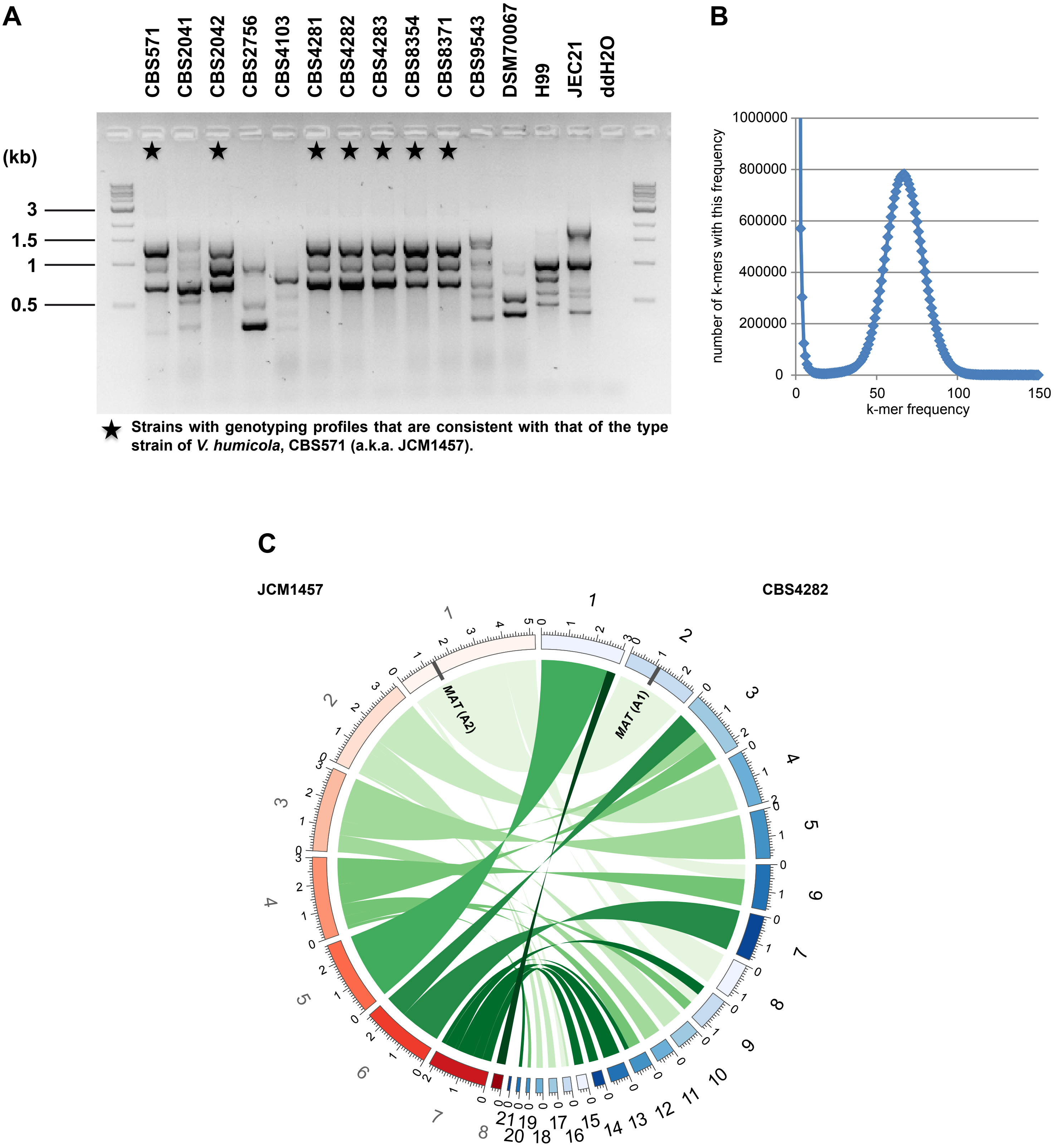


**S1 Figure.** Genome analysis of *V. humicola* strains. **A.** Genotyping of the *V. humicola* isolates with modified RAPD method. Stars highlight the isolates that show identical genotyping profiles as that of the *V. humicola* type strain, CBS571, and have ITS sequences that classify them as *V. humicola*. **B.** k-mer frequency distribution for *V. humicola* CBS4282 Illumina reads. A k-mer length of k=31 were used for the analysis. A single main peak can be observed. The rise in k-mer occurrence below a k-mer frequency of 10 is due to sequencing errors. The sum of k-mers from the main peak results in an estimate of 22.6 Mb for the size of the haploid genome. k-mer frequency analysis was performed as described in Traeger et al. (2013) [PLoS Genet 9: e1003820] and The Potato Genome Sequencing Consortium (2011) [Nature 475: 189–195]. **C.** Genome comparison between *V. humicola* strains JCM1457 (A2) and CBS4282 (A1). Regions of sequence similarity were determined by nucmer and plotted with Circos. Sizes are given in Mb. The positions of the *MAT* regions are indicated as grey bars.

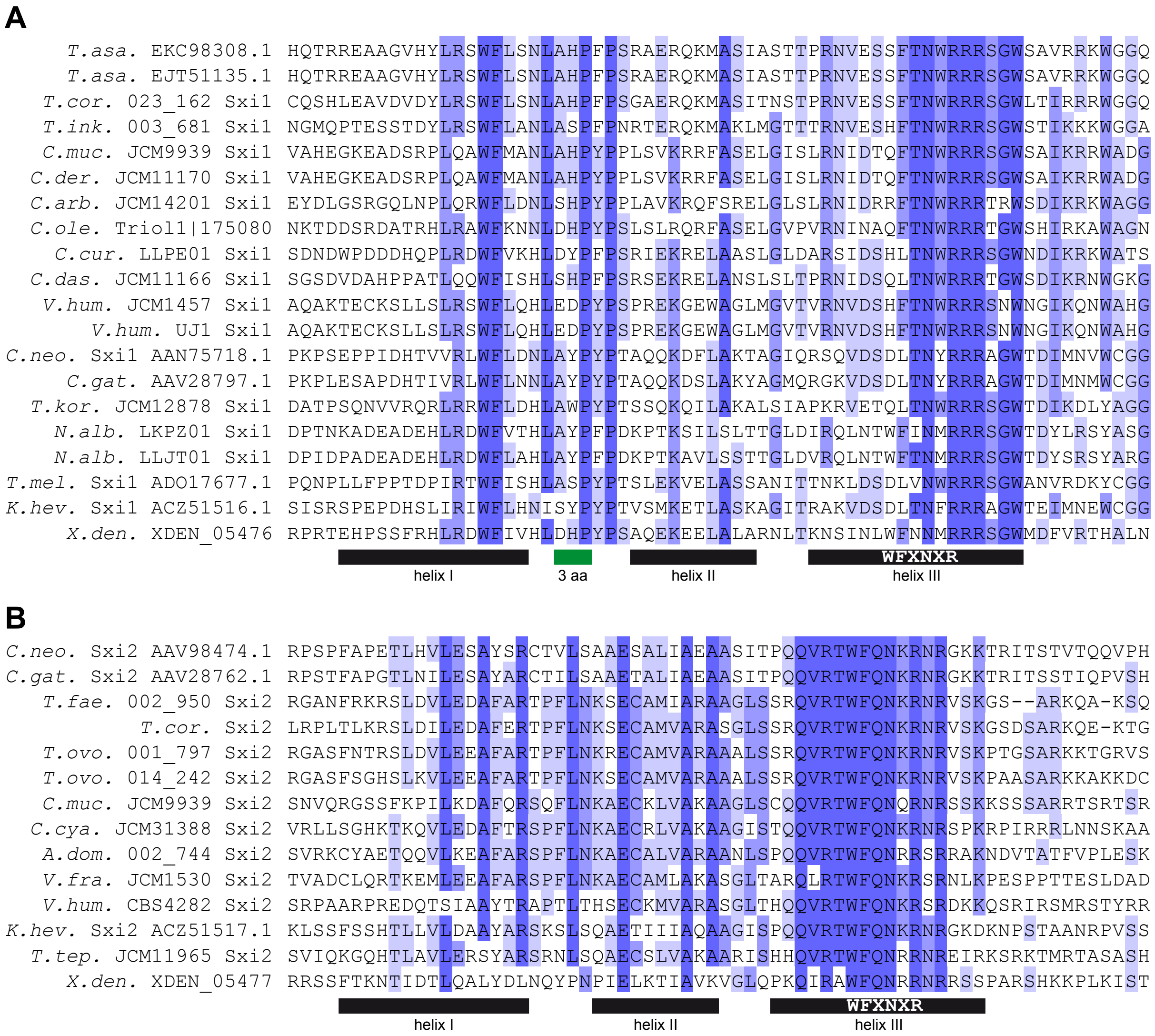


**S2 Figure.** Analysis of Sxi proteins from *Tremellomycetes*. **A.** Multiple alignment of the homeodomains from Sxi1 homologs (class HD1 homeodomain transcription factors) from *Tremellomycetes*. The three conserved helices are underlined in black. A three amino acid insertion characteristic for the HD1 homeodomain factors (in contrast to the HD2 homeodomains) is underlined in green. The conserved DNA binding motif WFXNXR within helix III is indicated. Labelling of helices and motifs according to Kües et al. (The Mycota XIV, 2011, p. 97-160. Springer, Berlin, Heidelberg). Species: *C.neo. Cryptococcus neoformans, C.gat. Cryptococcus gattii, C. arb. Cutaneotrichosporon arbiformis, C.cur. Cutaneotrichosporon curvatus, C. das Cutaneotrichosporon daszweskae, C. der Cutaneotrichosporon dermatis, C. muc. Cutaneotrichosporon mucoides, C.ole. Cutaneotrichosporon oleaginosum, K.hev. Kwoniella heveanensis, N.alb. Naganishia albida, T. kor. Takashimella koratensis, T.asa. Trichosporon asahii, T.cor. Trichosporon coremiiforme, T.ink. Trichosporon inkin, T.mel. Tremella mesenterica, V.hum. Vanrija humicola, X.den. Xanthophyllomyces dendrorhous*. **B.** Multiple alignment of the homeodomains from Sxi2 homologs (class HD2 homeodomain transcription factors) from *Tremellomycetes*. Labelling and species names as in A with addition of *A.dom. Apiotrichum domesticum, C. cya. Cutaneotrichosporon cyanovorans, T. tep. Takashimella tepidaria, T.fae. Trichosporon faecale, T.ovo. Trichosporon ovoides, V. fra. Vanrija fragicola*.

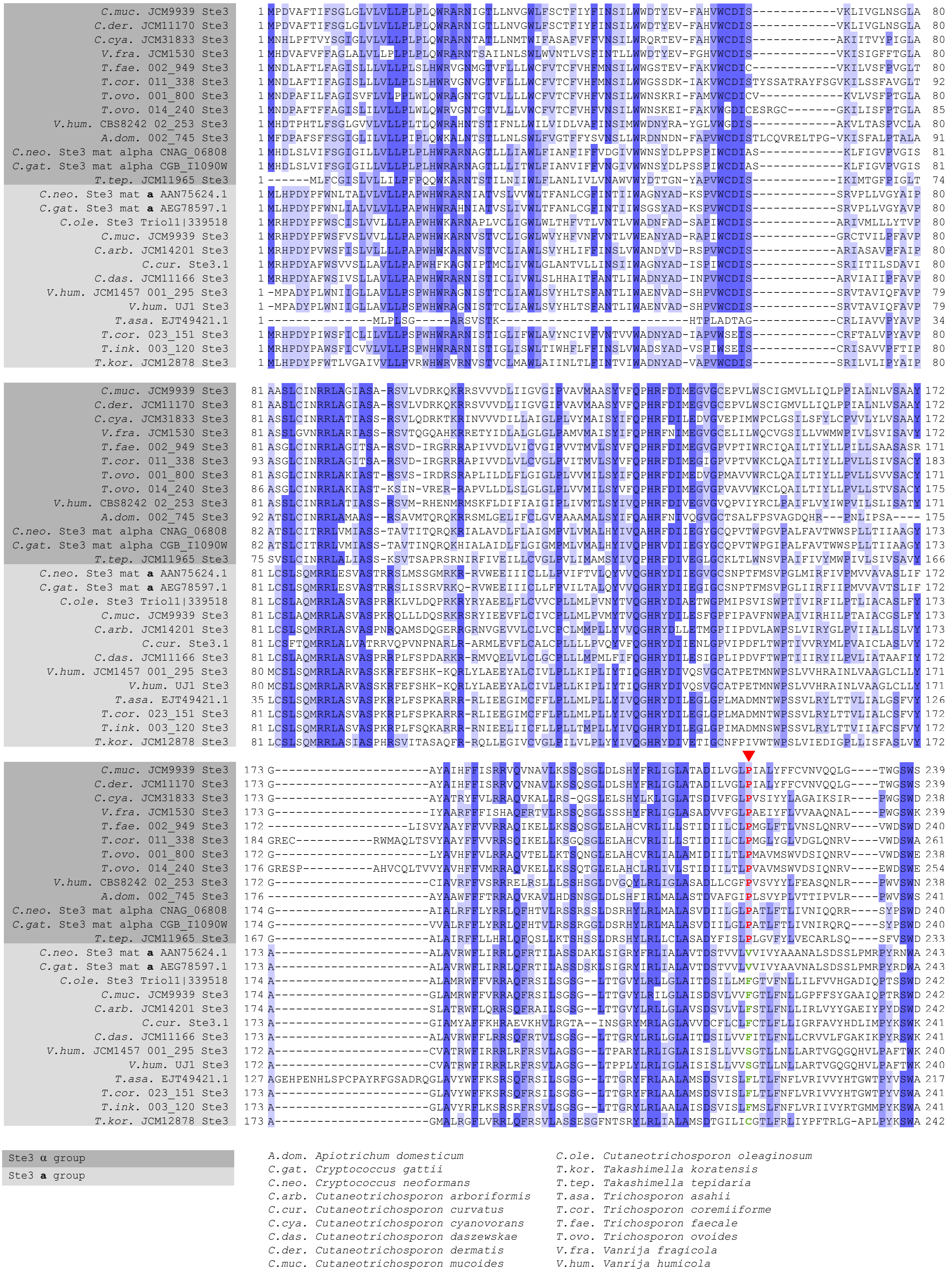


**S3 Figure.** Multiple alignment of the N-terminus of Ste3 homologs in the *MAT* loci of several *Tremellomycetes*. A conserved proline residue that is present in the α group proteins is indicated in red. It is not present in the **a**-specific Ste3 proteins from *Tremellomycetes*.

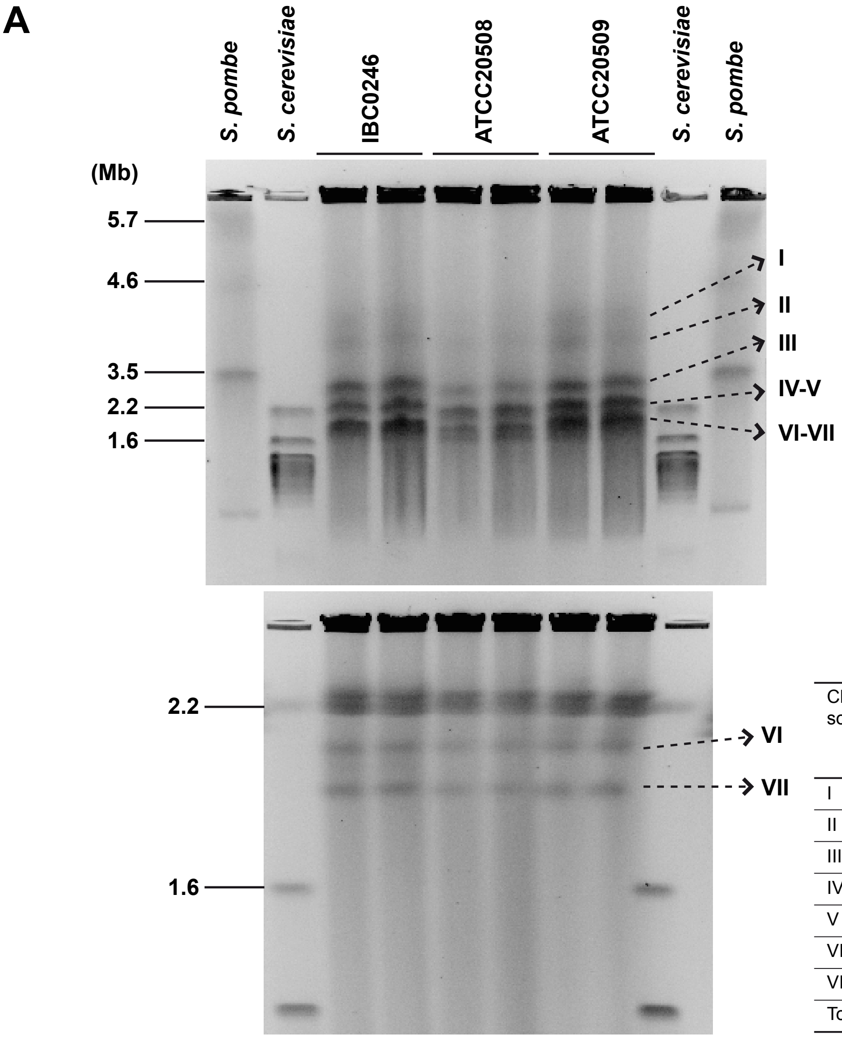


| Chromo-some | Size (~est., Mb) | Potentially corresponding contig in ATCC20508 assembly | Contig size (Mb) |
|---|---|---|---|
| I | 4 | 1 | 4.1 |
| II | 3.8 | 2 | 3.8 |
| III | 3 | 3 | 3.2 |
| IV | 2.3 | 4 | 2.7 |
| V | 2.3 | 5 | 2.2 |
| VI | 2 | 6 | 2.0 |
| VII | 1.8 | 7+8 | 0.9+0.8 |
| Total | 19.2 | | 19.8 |

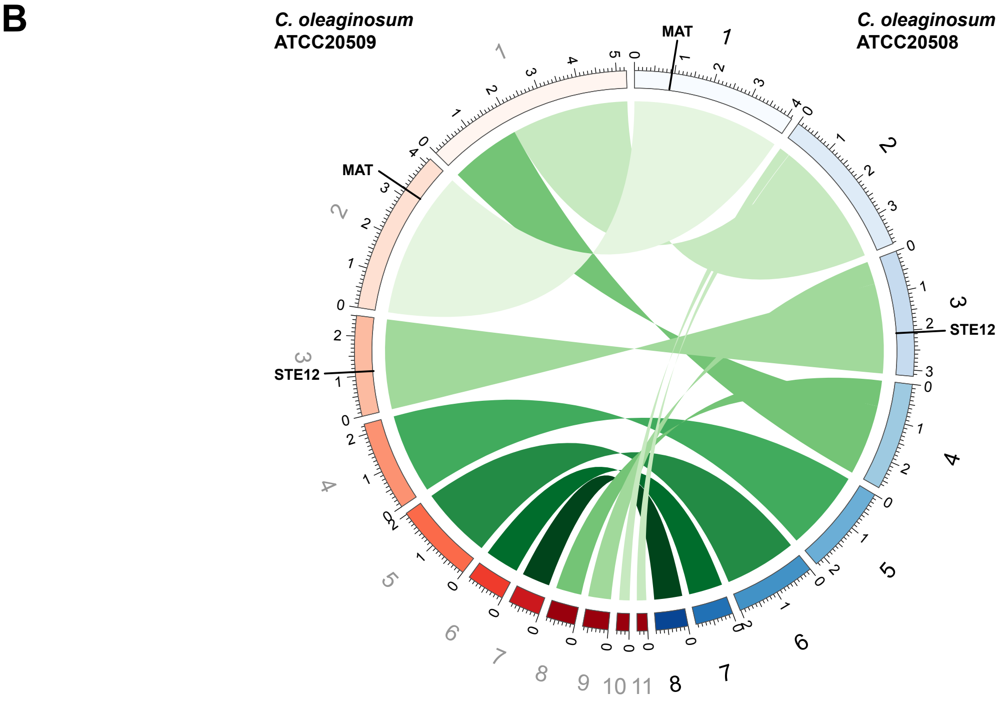


**S4 Figure.** Genome sequencing of *T. oleaginosus* ATCC20508 (*MAT* A2). The genome was sequenced with Pacific Biosciences SMRT sequencing and assembled into eight contigs with 8208 predicted genes. **A.** CHEF analysis of the three *C. oleaginosum* isolates. The image on top shows separation of the larger chromosomes, with *S. pombe* and *S. cerevisiae* serving as markers; the image at the bottom shows separation of the smaller chromosomes of the same three *C. oleaginosum* isolates, with only *S. cerevisiae* included as marker (please see Materials and Methods for details). Two plugs prepared from independent cultures were included for each isolate. The table summarizes the estimated sizes of the seven chromosomes identified by the CHEF analysis. The sizes of the six largest chromosomes correspond well to the sizes of the six largest contigs of the ATCC20508 assembly. Similar to the other two strains, ATCC20508 carries the *MAT* A2 allele and *STE12* on different contigs (contigs 1 and 3, respectively, see part B), making it unlikely that they are located on the same chromosome. **B.** Genome comparison between *C. oleaginosum* strains ATCC20508 (A2) and ATCC20509 (A2). Regions of sequence similarity were determined by nucmer and plotted with Circos. Sizes are given in Mb. The positions of the *MAT* regions and *STE12* are indicated.

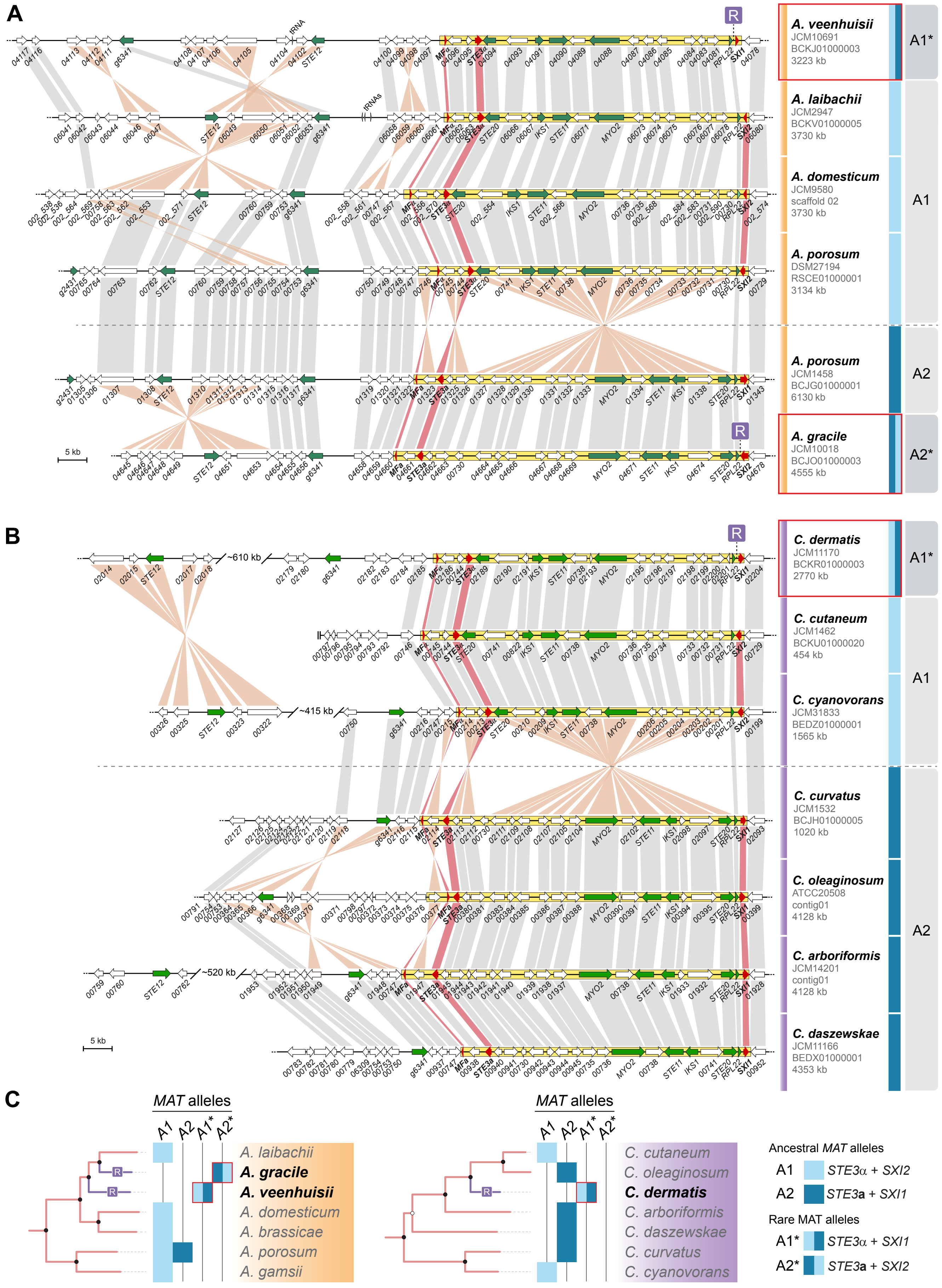


**S5 Figure.** *MAT* loci of *Apiotrichum* (**A**) and *Cutaneotrichosporon* (**B**) species. Classic mating type-defining genes (*SXI1* or *SXI2*, *STE3*, and the pheromone precursor genes) are shown in red, other genes that are part of the mating type locus or flanking the mating type locus (gene *g6341*) of *C. neoformans* are shown in green, and genes not present in the *C. neoformans* mating type locus are shown in white. The *MAT* allele is indicated on the right, with A1/A1* and A2/A2* *MAT* alleles shown above and below the dashed line, respectively. The proposed *MAT* locus region is enclosed in a yellow box in each strain. Orthologs are connected by grey or orange bars when in the same or opposite orientations, respectively. Strains carrying the A1* or A2* alleles are outlined in red. Predicted recent recombination events leading to the A1* and A2* alleles are indicated by the letter R. The hybrid species *C. mucoides* is not shown in this Figure, please refer to Fig S6 for an analysis including hybrid species. **C.** Phylogenetic trees showing species relationships within the genera *Apiotrichum* and *Cutaneotrichosporon*. The *MAT* alleles of the strains are indicated, and predicted recent recombination events leading to the A1* and A2* alleles are indicated by the letter R at the corresponding branches of the phylogenetic trees.

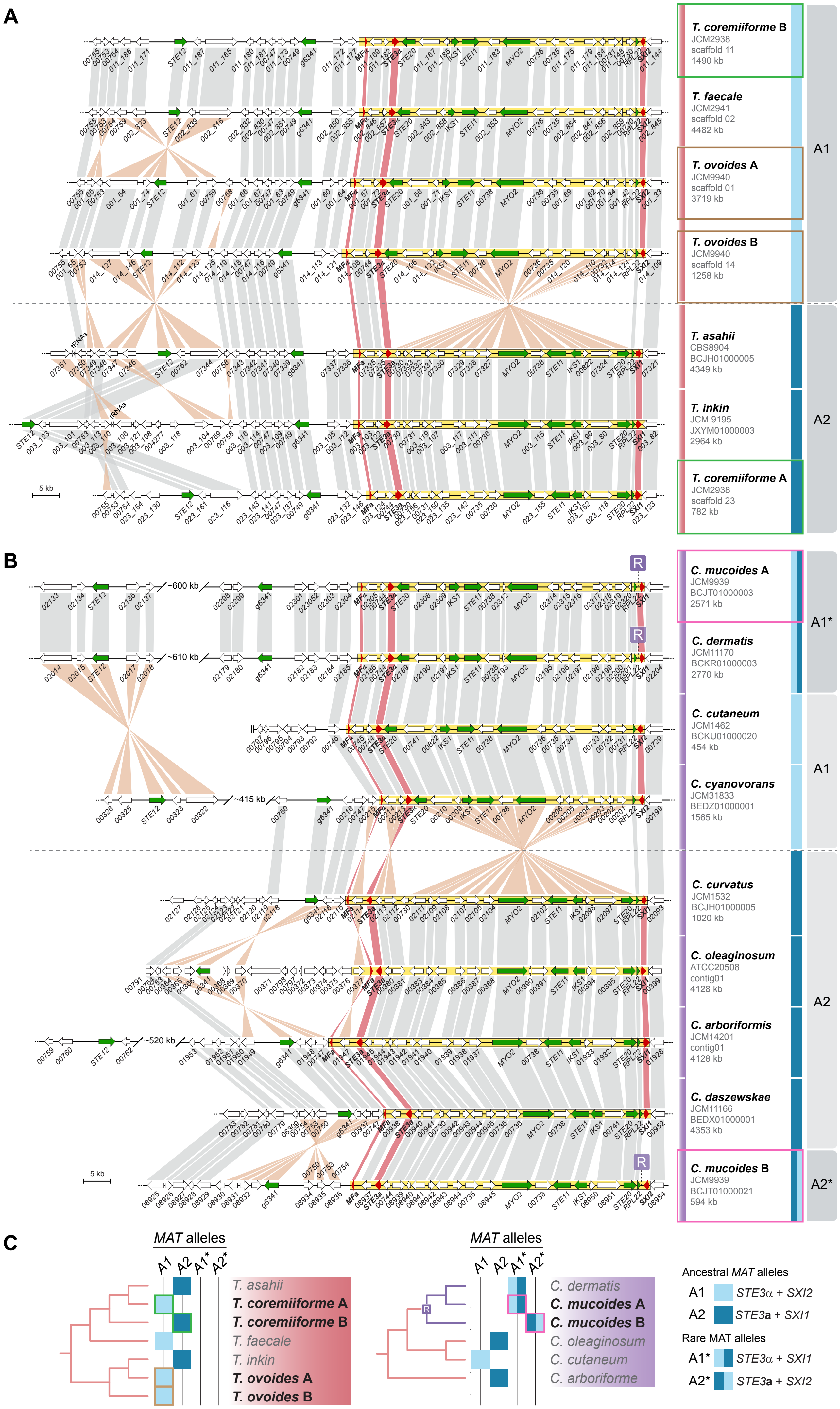


**S6 Figure.** *MAT* loci of *Trichosporonales* lineages containing hybrid species (*Trichosporon* species in **A**, *Cutaneotrichosporon* species in **B**). Classic mating type-defining genes (*SXI1* or *SXI2*, *STE3*, and the pheromone precursor genes) are shown in red, other genes that are part of the mating type locus or flanking the mating type locus (gene *g6341*) of *C. neoformans* are shown in green, and genes not present in the *C. neoformans* mating type locus are shown in white. The *MAT* allele is indicated on the right, with A1/A1* and A2/A2* *MAT* alleles shown above and below the dashed line, respectively. The proposed *MAT* locus region is enclosed in a yellow box in each strain. Orthologs are connected by grey or orange bars when in the same or opposite orientations, respectively. Predicted recent recombination events leading to the A1* and A2* alleles are indicated by the letter R. *MAT* alleles of hybrid strains are outlined by colored boxes around the strain names on the right. **C.** Phylogenetic trees showing species relationships within the genera *Trichosporon* and *Cutaneotrichosporon*. The *MAT* alleles of the strains are indicated, and a predicted recent recombination event leading to the A1* and A2* alleles in *Cutaneotrichosporon* is indicated by the letter R at the corresponding branch of the phylogenetic tree. Names of hybrid species are shown in bold. Each hybrid species carries two subgenomes designated A and B, with homeologs for each gene still present in both subgenomes [Genome Res. (2016) 26:1081-90; Yeast (2018) 35:99-111]. For more information about the hybrid strains, please see Text S2.

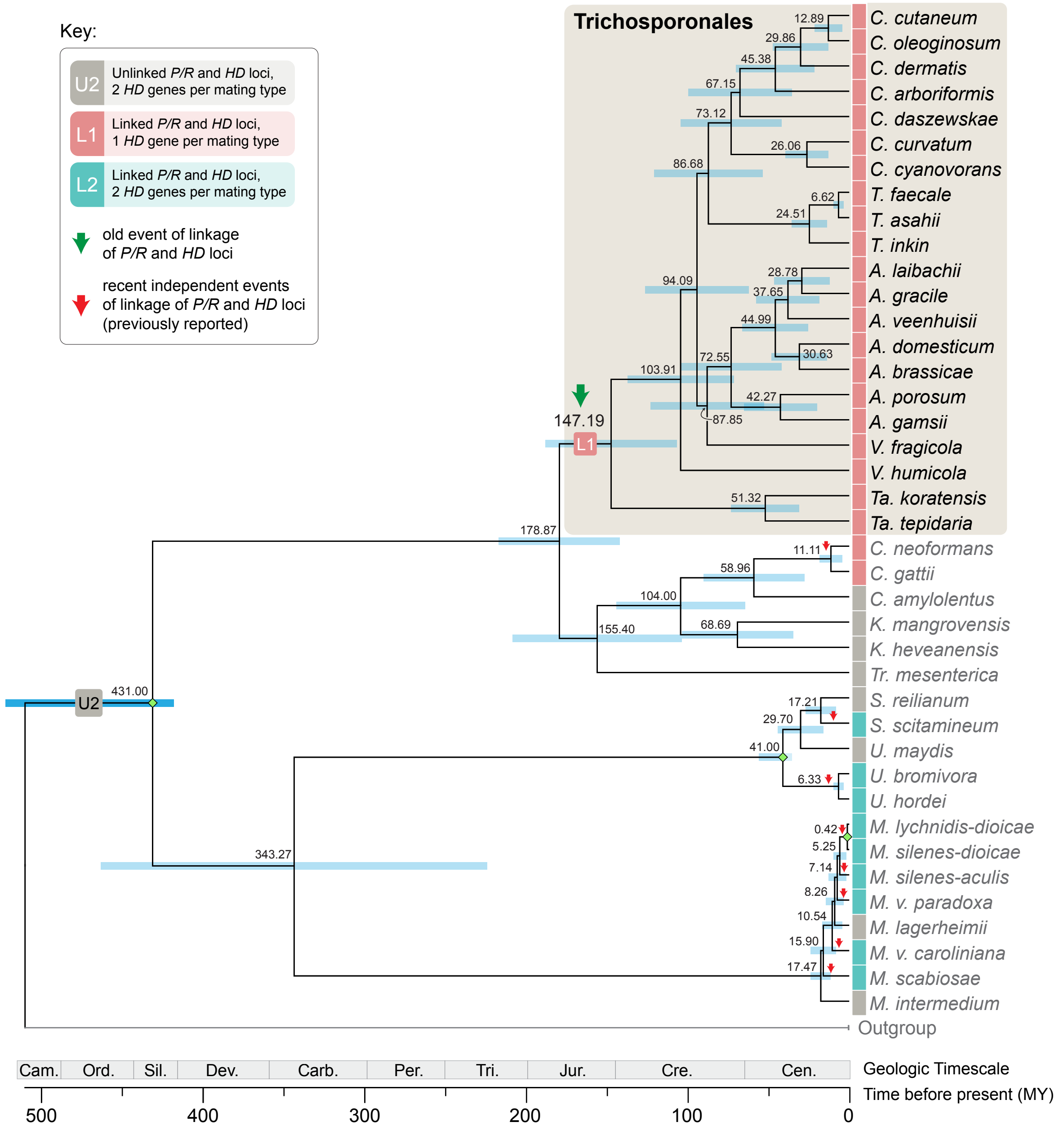


**S7 Figure.** Analysis of divergence times of basidiomycete lineages. The reconstructed species tree shown in Fig 1, with branch lengths in the units of number of substitutions per site, was used as input and transformed into an ultrametric tree with relative times. The final timetree was obtained by converting the relative node ages into absolute dates by using three calibration constraints: 0.42 million year (MY) corresponding to the divergence between *Microbotryum lychnidis-dioicae* and *Microbotryum silenes-dioicae* [Mol Biol Evol. (2011) 28:459-71]; 41 MY for the *Ustilago* - *Sporisorium* split; and 413 MY representing the minimum age of Basidiomycota. The latter two calibration points were obtained from the Timetree website (http://www.timetree.org/), which should be referred to for additional information and references. Numbers on tree nodes indicate the inferred dates of speciation. Events of mating-type loci linkage are indicated by red or green arrows (see key). The blue bars correspond to 95% confidence intervals.

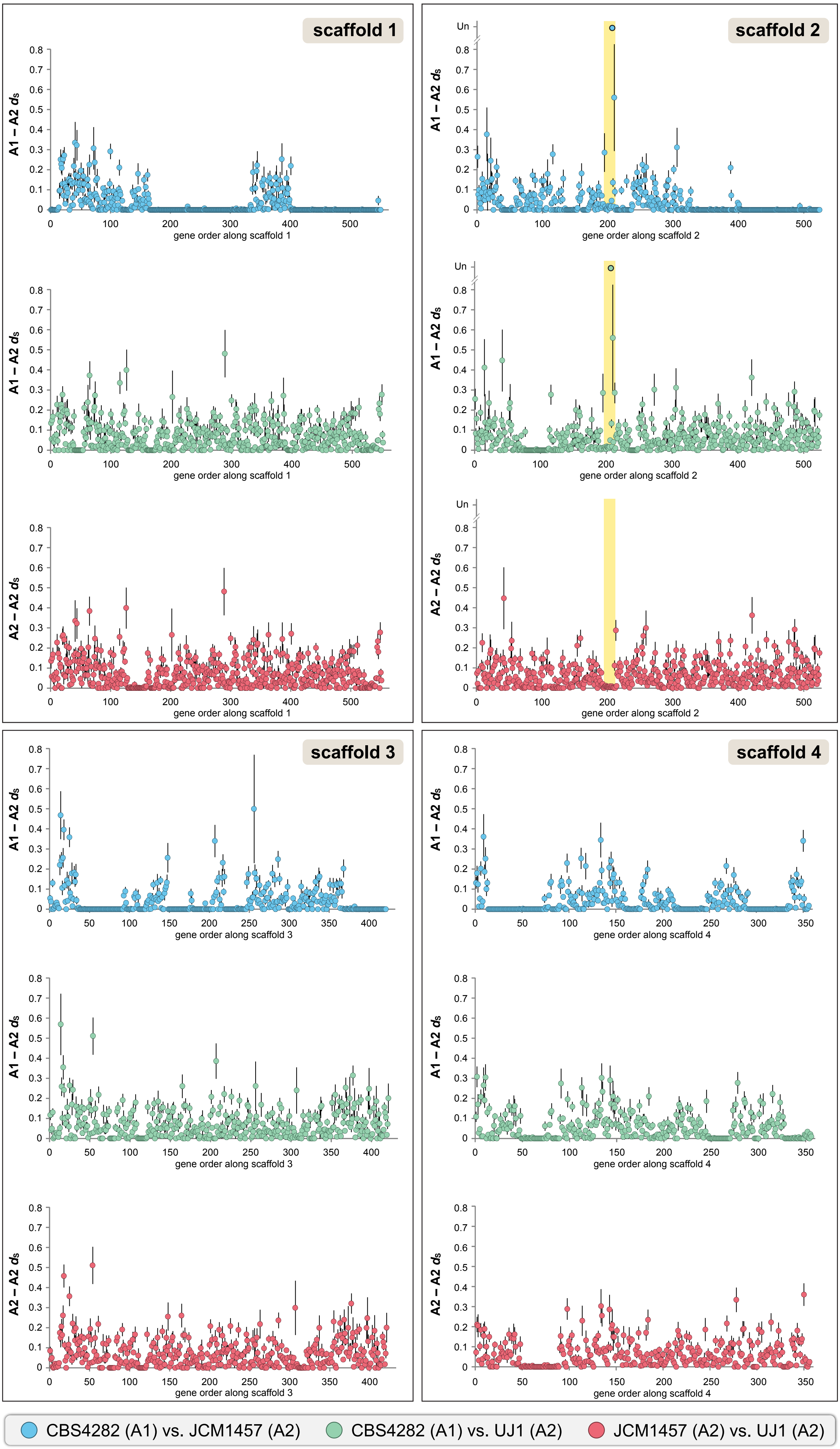


**S8 Figure.** Synonymous divergence between alleles in different *V. humicola* strains. Synonymous substitutions per synonymous site and standard errors ($d_S$ +/- SE) are shown for pairwise comparisons between strains CBS4282, JCM1457, and UJ1 for genes on the four longest scaffolds (scaffold 2 contains the *MAT* locus) of strain CBS4282. Only genes for which a $d_S$ value could be calculated are shown except for the *STE3* genes, for which no $d_S$ value could be calculated in comparisons of strains with different mating types, and which are represented at the value Un (undetermined). The location of the core *MAT* region (between the *HD* and *P/R* genes) on scaffold 2 of CBS4282 is labelled in yellow.

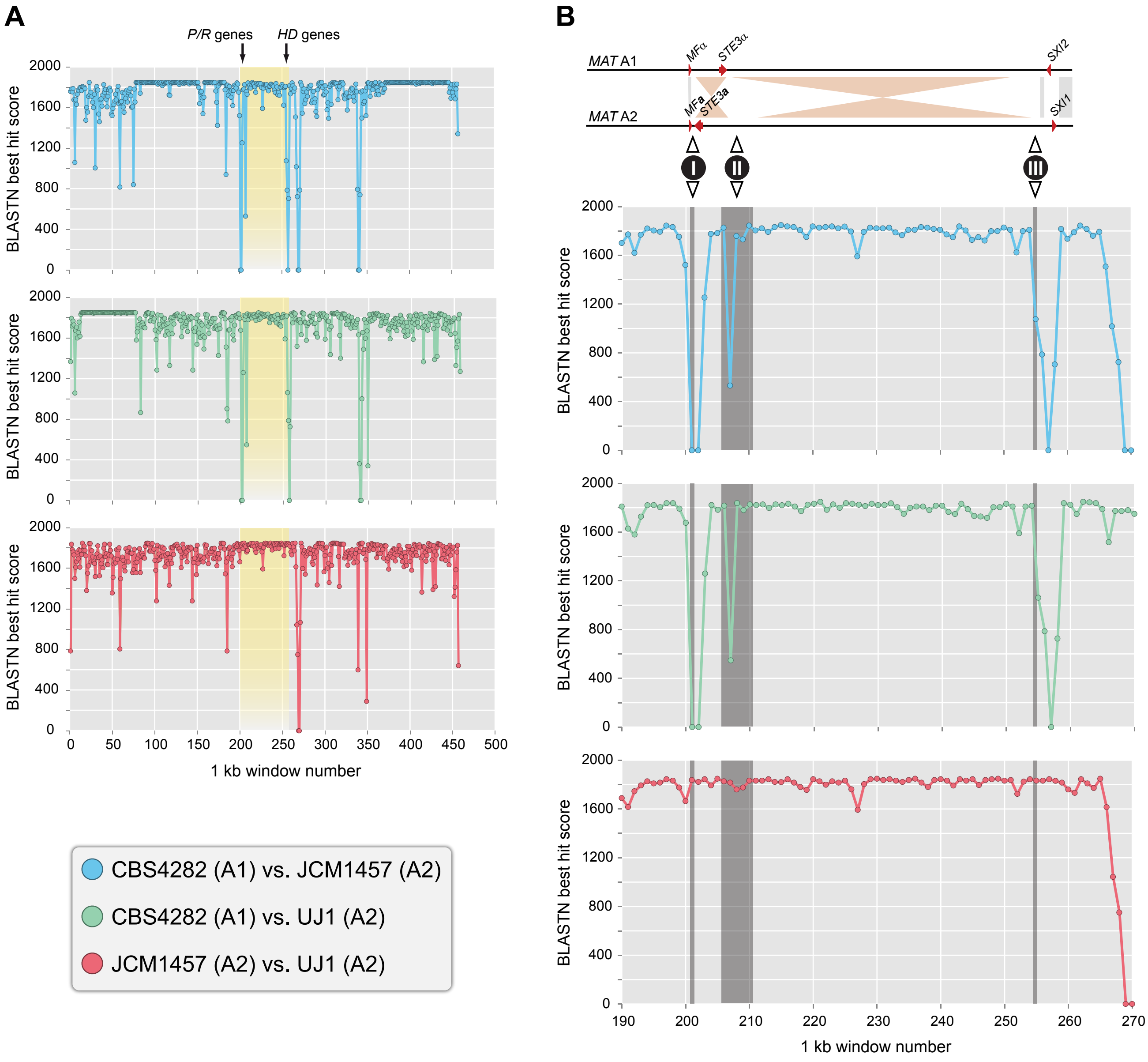


**S9 Figure.** BLASTN scores of sliding window comparisons of *MAT* regions from *V. humicola* strains. The *MAT* loci (region between the pheromone gene/*STE3* and the corresponding HD gene) and 200 kb upstream and downstream from strains JCM1457 and UJ1 were split into fragments of 1 kb and used in BLASTN comparisons against the corresponding regions from the indicated strains. BLASTN scores for the best hits were plotted against the corresponding fragment. Higher BLASTN scores indicate higher sequence similarity, more diverse regions have lower scores. **A.** Overview for the three pairwise comparisons. The region shaded in yellow is the core *MAT* region (between the *P/R* genes and the corresponding *HD* gene). **B.** Detail from (A) showing 1 kb windows #190-270, which encompass the *MAT* region. Grey shading indicates the windows that harbor potential inversion breakpoints for two inverted regions that are indicated in a schematic view above. The three regions (**I-III**) have a lower degree of similarity in comparisons of A1 and A2 strains than in the comparison of the two A2 strains (UJ1 and JCM1457).

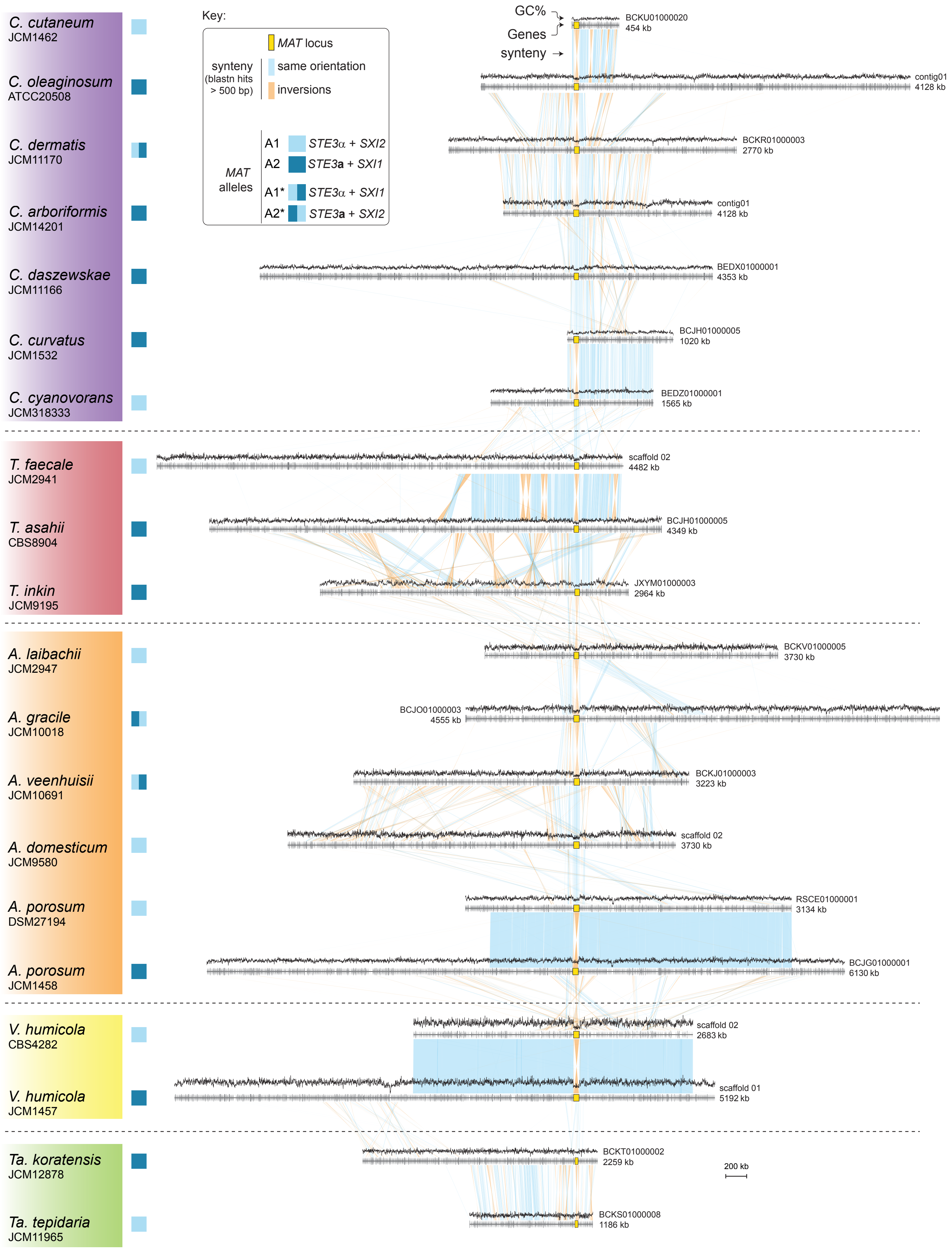


**S10 Figure.** Synteny analysis of *MAT*-containing scaffolds. Linear synteny comparison along the *MAT*-containing scaffolds was generated with Easifig [Bioinf. (2011) 27:1009-10] using a minimum length of 500 for BLASTN hits to be drawn.

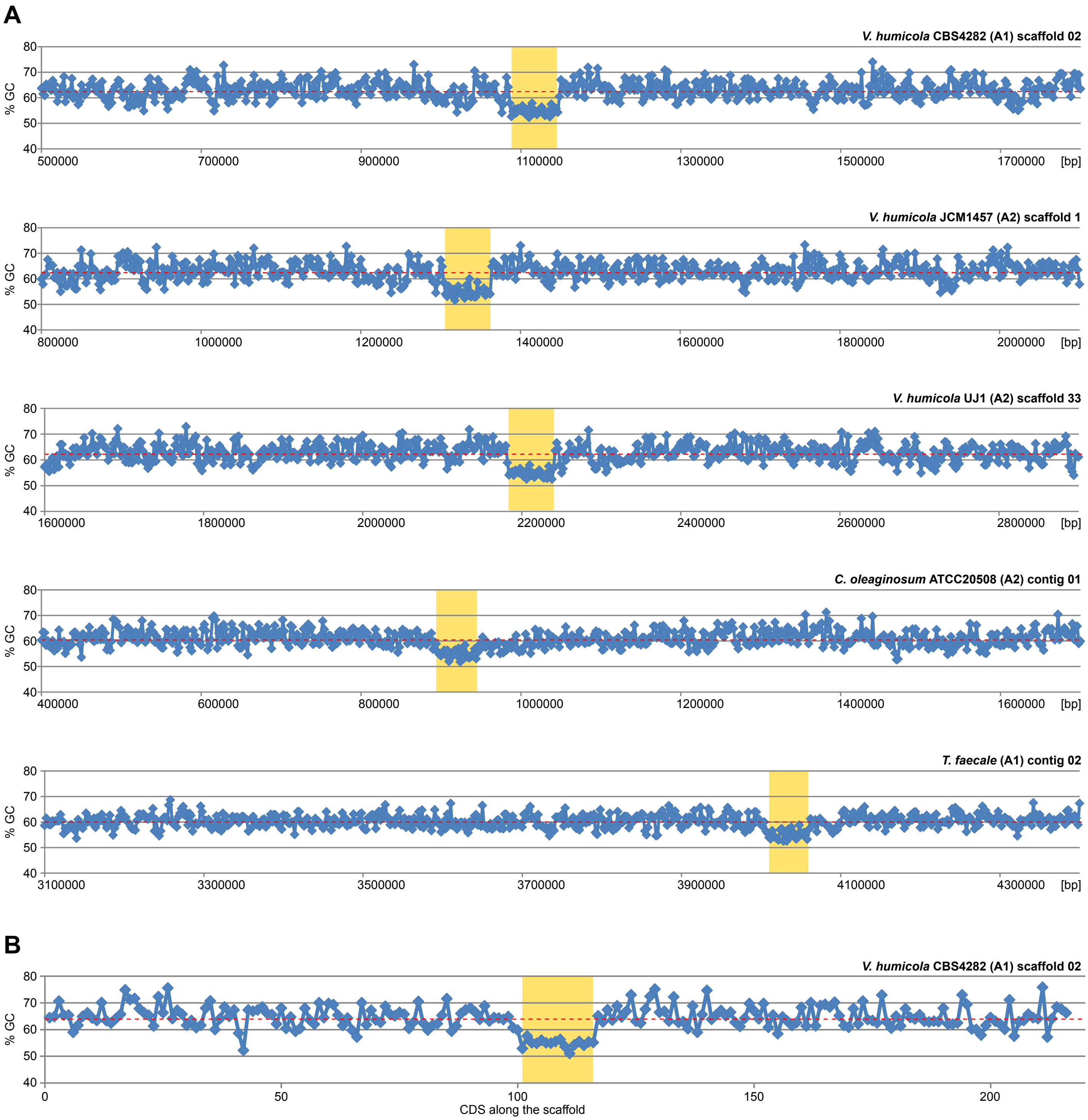


**S11 Figure.** GC content of the *MAT* regions and surrounding regions in several *Trichosporonales* strains. **A.** GC content in percent is plotted against the scaffold coordinates for the *MAT*-containing scaffolds. GC content was determined in windows of 2000 bp along each scaffold. The core *MAT* regions (between *P/R* and the *SXI* genes) are shaded in yellow for each scaffold. The average genomic GC content for each strain is indicated by a dashed red line. GC content in the core *MAT* region is lower than in the surrounding regions, which have GC contents that vary around the average GC content. **B.** GC content was calculated for each coding sequence (CDS) and plotted for 100 genes upstream and downstream from the core *MAT* region of strain *V. humicola* CBS4282.The average GC content for CDSs of this strain is indicated by a dashed red line, the core *MAT* region is shaded in yellow. The lower GC content observed in the genomic sequence of the *MAT* region is also observed when analyzing only CDS sequences.

**S1 Text. Pheromone genes of *Tremellomycetes*.**

One difference between the *Trichosporonales* and several *Tremellales MAT* loci is the presence of only one pheromone precursor gene in the analyzed *Trichosporonales MAT* loci (Fig 2), whereas more than one pheromone precursor gene is present within the *P/R* locus in the tetrapolar *Tremellales* species *C. amylolentus*, and *T. wingfieldii* as well as in the bipolar species *C. neoformans* and *Cryptococcus gattii* [1-4]. In all cases, the pheromone precursor genes can be found in the vicinity of the pheromone receptor gene *STE3*. The *STE3*α-associated pheromones are slightly shorter in both the *Trichosporonales* (26-42 amino acids) as well as the *Tremellales* (38 amino acids) than their *STE3***a**-associated counterparts (27-47 in the *Trichosporonales*, and 39-42 amino acids in the *Tremellales*) (see Figure below). The predicted *Trichosporonales* pheromones have the characteristic C-terminal CAAX motif of lipopeptide pheromones, where A is an aliphatic amino acid [5]. However, the *Trichosporonales* pheromones have a shorter N-terminus compared to the *Tremellales* pheromones, except for those from the *Vanrija* species, *C. curvatus* and *C. cyanovorans* (see Figure below).

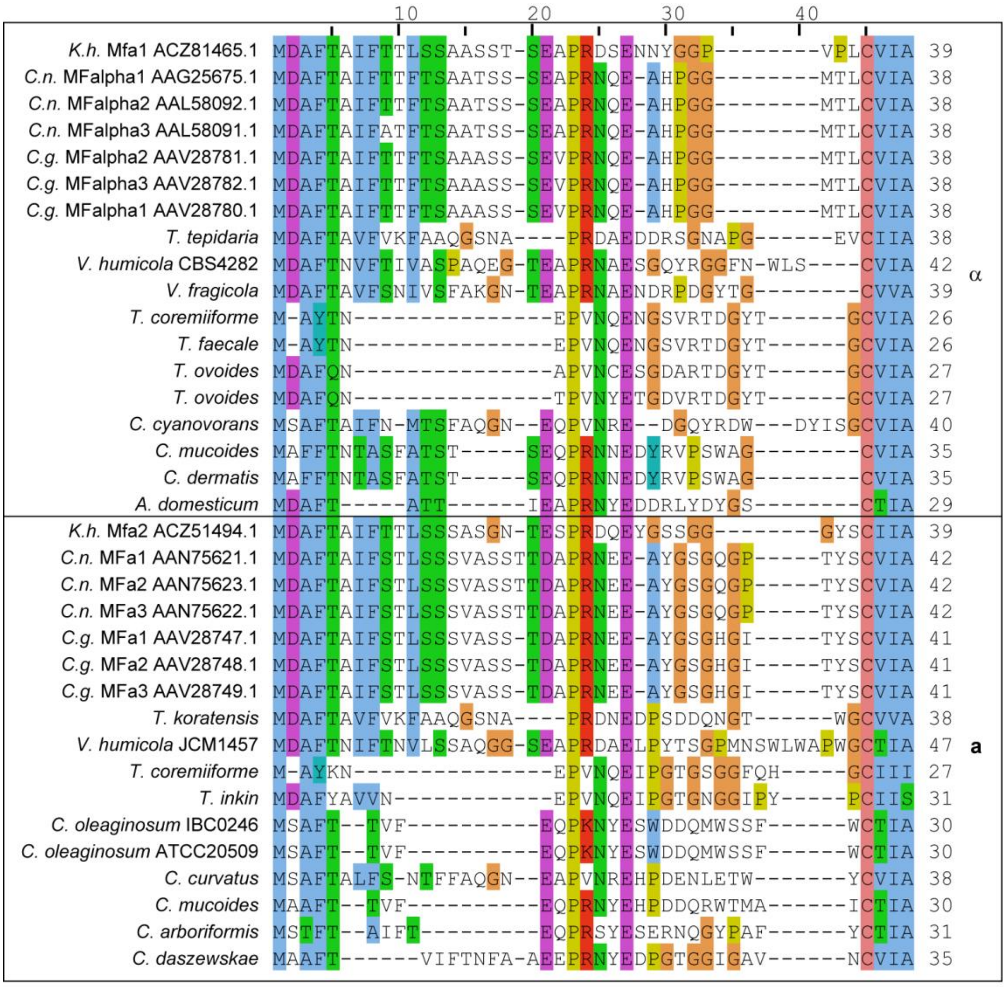


**Figure for Text S1.** Multiple alignment of (putative) pheromones from *Tremellomycetes*. The pheromones associated with the *STE3*α receptor gene are given on top, pheromones associated with the *STE3***a** receptor gene below. For *Tremellales* sequences, accession numbers are given after the species abbreviations (*C.n.*, *Cryptococcus neoformans*; *C.g.*, *Cryptococcus gattii*; *K.h.*, *Kwoniella heveanensis*), *Trichosporonales* sequences were identified in this study.

**S2 Text. *MAT* loci in hybrid species**

Among the sequenced *Trichosporonales* genomes are three genomes derived from the hybrid species *T. coremiiforme*, *T. ovoides*, and *C. mucoides*. These species are most likely the result of prior inter-species hybridizations between closely related *Trichosporon* or *Cutaneotrichosporon* species that occurred during the evolution of the *Trichosporonales* lineage [1, 2]. Each hybrid genome has approximately twice the size of the non-hybrid genomes and still contains homeologs for the majority of genes [1, 2]. This includes the *MAT* locus, and thus, there are two *MAT* loci, each with fused *HD* and *P/R* loci, present in each genome originating from the two different species that underwent hybridization (Fig S5). The analyzed genome of the hybrid species *C. mucoides* carries both the A1* as well as the A2* allele (Fig S5B). Its closest haploid relative is *C. dermatis* [2], which carries the A1* allele, making it likely that any recombination leading to the A1* and A2* alleles occurred in the ancestor of the lineage leading to *C. mucoides* and *C. dermatis*, because the other investigated *Cutaneotrichosporon* strains from different species carry the A1 or A2 alleles. An alternative hypothesis could be that the A1* and A2* alleles are the products of an independent loss of one of the two *HD* genes.

Hybridization occurred also in the *Trichosporon* lineage [1], and the two hybrid species *Trichosporon coremiiforme* and *Trichosporon ovoides* also carry two *MAT* loci (each a fusion of one *HD* and one *P/R* locus) per genome. The analyzed *T. coremiiforme* genome contains an A1 and an A2 allele, while the analyzed *T. ovoides* genome carries two A1 alleles (Fig S5A). Sequence similarity between the two alleles present in each of the three hybrid strains, which represent three different hybrid species, is much lower than between the two alleles present in the *V. humicola* strains CBS4282 and JCM1457 (see Figure below). Thus, it is likely that the two alleles in each of the hybrid strains originated from the different parental species forming the hybrid. One question is why the presence of two different *MAT* alleles, and consequently an *SXI1* and *SXI2* gene, in *C. mucoides* and *T. coremiiforme* does not initiate sexual development including meiosis. One possibility is that the HD proteins from different species are not able to form a functional heterodimer. Alternatively, genetic events, e.g. loss of homeologous genes, might have occurred following fusion that prevent self-fertile development. Another interesting point is the finding of two A1 alleles in the analyzed *T. ovoides* genome (Fig S5A), making it unlikely that the fusion of the parental strains was part of the cell fusion that occurs during mating of strains carrying different alleles during sexual reproduction. It is possible that this was a fusion of vegetative cells, or part of a "unisexual reproduction" (same-sex mating) life cycle in the parental species. Another possibility could be mating between a diploid A1/A2 hybrid with an A1 haploid, or between two A1/A2 hybrids. In these cases, random assortments will lead to progeny with two same *MAT* alleles.


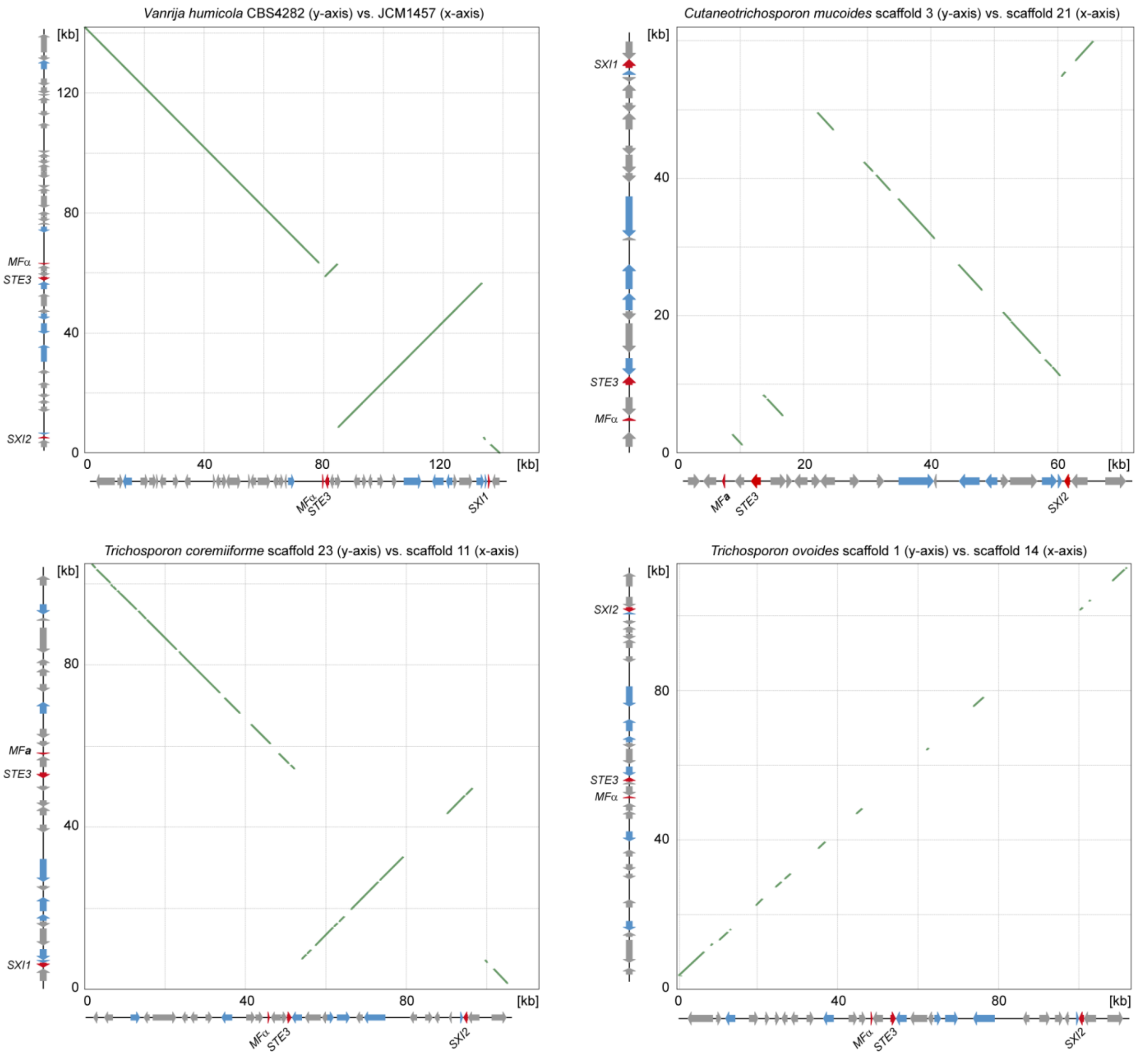


**Figure for Text S2.** Dot plots of sequence comparisons between *MAT* loci. Nucleic acid sequences of *MAT* loci for two *V. humicola* strains carrying the A1 and A2 alleles, and between the two alleles present in the hybrid strains *C. mucoides*, *T. coremiiforme*, and *T. ovoides* were performed with the nucmer algorithm from the MUMmer package. Dot plots were generated with gnuplot. The highest degree of similarity is observed between the two V. humicola strains, whereas the two alleles within each of the three hybrid strains are less similar to each other confirming the inter-species hybridization origin of the strains. The inversion of the region between the *STE3* and *SXI* genes can be seen in comparisons of different alleles (in the *V. humicola*, *C. mucoides*, and *T. coremiiforme* comparisons).
